# A toolbox for ablating excitatory and inhibitory synapses

**DOI:** 10.1101/2024.09.23.614589

**Authors:** Aida Bareghamyan, Changfeng Deng, Sarah Daoudi, Shubash C. Yadav, Xiaocen Lu, Wei Zhang, Robert E. Campbell, Richard H. Kramer, David M. Chenoweth, Don B. Arnold

**Affiliations:** Department of Biology, Division of Molecular and Computational Biology, University of Southern California, Los Angeles, CA 90089, USA; Neuroscience Graduate Program, University of Southern California, Los Angeles, CA 90089, USA; Department of Chemistry, School of Arts and Sciences, University of Pennsylvania, Philadelphia, PA 19104, USA; Department of Molecular and Cell Biology, University of California, Berkeley, California, 94720, USA; Department of Chemistry, Faculty of Science, University of Alberta, Edmonton, Alberta, T6G 2G2, Canada; Department of Chemistry, Graduate School of Science, The University of Tokyo, Bunkyo-ku, Tokyo, 113-0033, Japan; Department of Biomedical Engineering, Viterbi School of Engineering, University of Southern California, Los Angeles, CA 90089, USA

**Keywords:** PSD-95, Gephyrin, synapse, ablation, E3 ligase, photoactivatable

## Abstract

Recombinant optogenetic and chemogenetic proteins are potent tools for manipulating neuronal activity and controlling neural circuit function. However, there are few analogous tools for manipulating the structure of neural circuits. Here, we introduce three rationally designed genetically encoded tools that use E3 ligase-dependent mechanisms to trigger the degradation of synaptic scaffolding proteins, leading to functional ablation of synapses. First, we developed a constitutive excitatory synapse ablator, PFE3, analogous to the inhibitory synapse ablator GFE3. PFE3 targets the RING domain of the E3 ligase Mdm2 and the proteasome-interacting region of Protocadherin 10 to the scaffolding protein PSD-95, leading to efficient ablation of excitatory synapses. In addition, we developed a light-inducible version of GFE3, paGFE3, using a novel photoactivatable complex based on the photocleavable protein PhoCl2c. paGFE3 degrades Gephyrin and ablates inhibitory synapses in response to 400 nm light. Finally, we developed a chemically inducible version of GFE3, chGFE3, which degrades inhibitory synapses when combined with the bio-orthogonal dimerizer HaloTag ligand-trimethoprim. Each tool is specific, reversible, and capable of breaking neural circuits at precise locations.

## Introduction

Neural circuits, groups of neurons connected by synapses, are the basic units of computation in the brain and are fundamental to understanding its function and pathology. Although there are many genetically encoded tools for modulating neuronal function, existing tools for eliminating synapses are inadequate for providing a clear understanding of neural circuits. Previously, we developed GFE3, a protein that can ablate inhibitory synapses efficiently, rapidly, and without toxicity by targeting an E3 ligase to the scaffolding protein Gephyrin^1^. GFE3 consists of GPHN.FingR (**F**ibronectin **in**trabody **g**enerated by m**R**NA display), a recombinant, antibody-like protein, which binds Gephyrin at inhibitory synapses^2,3^ fused to the RING domain of the E3 ligase XIAP (RING_XIAP_) ^4^. GFE3 mediates the ubiquitination of Gephyrin and disassembly of inhibitory synapses. Importantly, it specifically targets postsynaptic sites containing GABA_A_ receptors and can be expressed in genetically determined cell types. Thus, GFE3 can manipulate circuits in a manner that is not possible with optogenetics^5^, DREADDs^6^, and modulators of neurotransmitter release^7^, which don’t target specific postsynaptic receptors, or with traditional pharmacological approaches that target specific receptors but do not allow for cell-type specificity. GFE3 has been used to probe the contribution of inhibitory inputs to LTP in hippocampal circuits^8^, recurrent loops in the oscillator that drives rhythmic whisking^9,10^, and the temporal coordination of vocalization and inspiration^11^. However, no tool analogous to GFE3 that ablates excitatory synapses currently exists. Furthermore, because GFE3 is constitutively active, its temporal and spatial resolution is limited.

In this study, we generated three tools for ablating synapses based on GFE3: 1. PFE3, an excitatory synapse ablator; 2. paGFE3, a photoactivatable version of GFE3; 3. chGFE3, a chemically activated version of GFE3. We show that the expression of PFE3 in cultured neurons causes the loss of PSD-95 puncta and excitatory synapses, which is reversible. In vivo, its expression leads to the functional loss of excitatory synaptic transmission. We generated paGFE3 by incorporating GPHN.FingR and the RING_XIAP_ into a novel photoactivatable complex based on the photocleavable protein PhoCl2c^12,13^. paGE3 is activated by 400 nm light and causes ablation of inhibitory synapses within 5 hr of exposure that is subsequently reversible. In the absence of 400 nm light, paGFE3 has no background activity. In addition, the expression of paGFE3 labels inhibitory synapses, allowing their size and location to be monitored before and after ablation. chGFE3, a chemogenetic version of GFE3 analogous to paGFE3, mediates efficient, reversible degradation of labeled synapses when a cell-permeant chemical is added.

## Results

### Degrading exogenous PSD-95 through ubiquitination

Initially, to generate an excitatory synapse ablator, we fused RING_XIAP_ to PSD-95.FingR, which binds at high affinity and specificity to the scaffolding protein PSD-95, which is found at excitatory synapses^14^. However, PSD-95.FingR-RING_XIAP_ did not efficiently ablate PSD-95 (data not shown). As an alternative, we fused PSD-95.FingR to the RING domain of the E3 ligase Mdm2 (RING_Mdm2_), which is necessary for PSD-95 degradation and excitatory synapse ablation during homeostatic plasticity^15^. To determine whether PSD-95.FingR-RING_Mdm2_ degrades PSD-95, we examined how co-expression with PSD-95.FingR-RING_Mdm2_ affected the expression of exogenous PSD-95-myc in COS7 cells. Cells transfected with PSD-95 alone had a PSD-95-myc expression level of 203 ± 32 au, as measured by western blot, while the expression level of PSD-95 in cells co-expressing PSD-95-myc and PSD-95.FingR-RING_Mdm2_ reduced to 107 ± 12 au, a ∼50% decrease (n = 5, p < 0.05, ANOVA with multiple comparisons). To determine whether this reduction in PSD-95 was due to ubiquitination, we tried a third condition, where the ubiquitination inhibitor TAK243^16^ was added to cells co-expressing PSD-95-myc and PSD-95.FingR-RING_Mdm2._ In this case, PSD-95 expression levels increased to 239 ± 31 au, a ∼20% increase relative to cells expressing PSD-95-myc alone, which was not significant (n = 5, p > 0.6, ANOVA with multiple comparisons). Furthermore, cells co-expressing PSD-95-myc and PSD-95.FingR-RING_Mdm2_ without TAK243 had ∼55% less PSD-95 vs. those with TAK243, a significant difference (p < 0.05, ANOVA with multiple comparisons, **Fig. 1A, B, S1B**).

**Figure 1:**
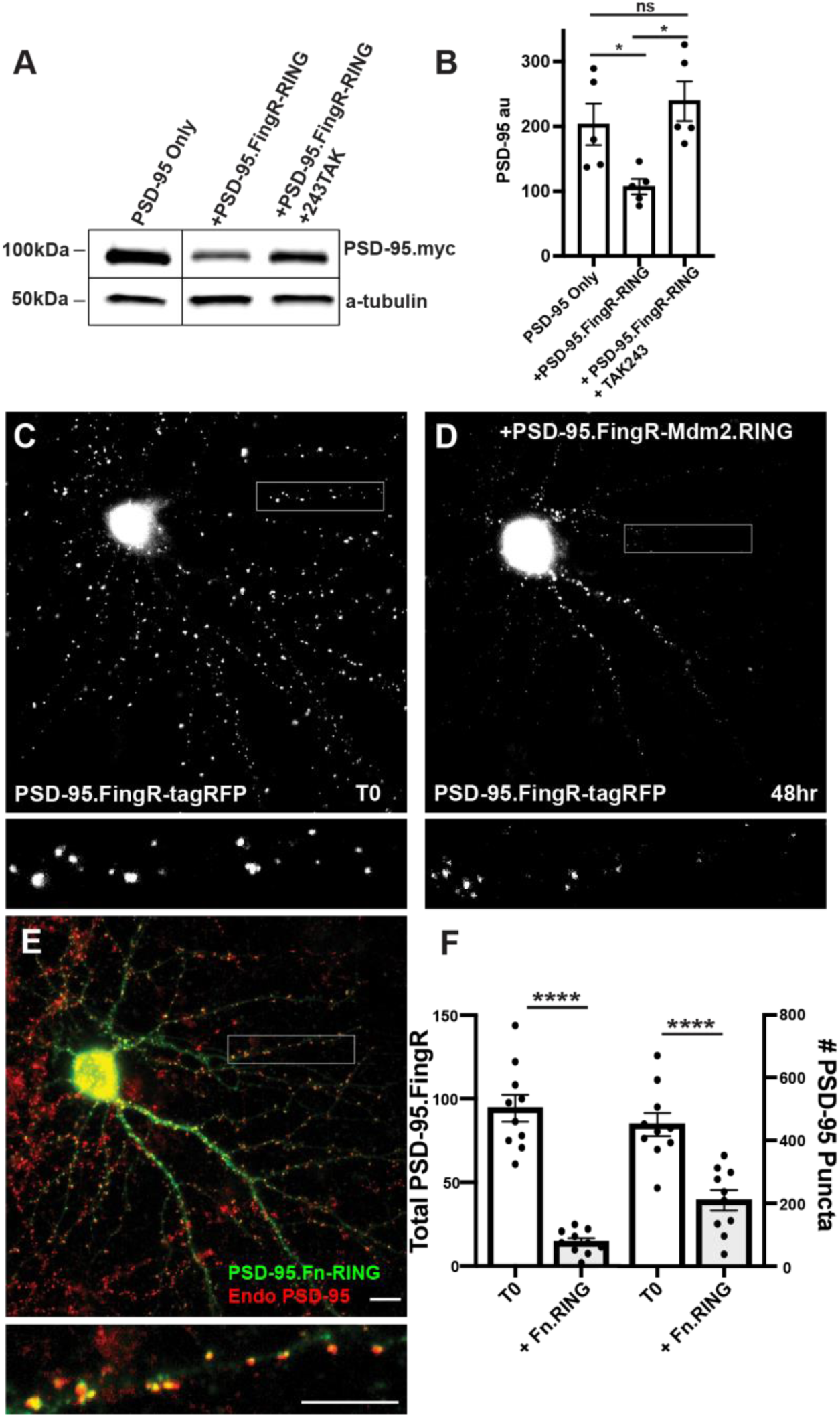
Mdm2.RING ubiquitinates PSD-95. **A)** Comparison of cell lysate from COS7 cells transfected with PSD-95-myc alone, PSD-95-myc + Dox-induced TRE-PSD-95.FingR-RING_MDM2_, and PSD-95-myc + PSD-95.FingR-RING _MDM2_ + TAK243, a ubiquitination inhibitor. Cells expressing TRE-PSD-95.FingR-RING_MDM2_ were treated with 1μg/ml Doxycycline for 4 hr to induce expression of PSD-95.FingR-RING and showed a reduction in PSD-95-myc. Cells treated with 20 μM TAK243 showed no apparent reduction in PSD-95-myc. **B)** Quantitation showed a significant reduction in PSD-95 expression when co-expressed with PSD-95.FingR-RING_MDM2_ in COS7 cells, but not when PSD-95 is expressed alone or when PSD-95 and PSD-95.Fn-RING _MDM2_ are co-expressed with 20 μM TAK243. * p < 0.05, ANOVA with multiple comparisons. ns, p > 0.05. **C)** Cultured cortical neuron expressing transcriptionally regulated PSD-95.FingR-tagRFP before induction of PSD-95.FingR-HA-RING_Mdm2_ expression with Dox. **D)** Same neuron as in C) after induction of PSD-95.FingR-HA-RING_MDM2_ expression with Dox shows a reduction in PSD-95.FingR-tagRFP labeling. **E)** Immunostaining of the neuron in D) for PSD-95.FingR-HA-RING_MDM2_ (green) and endogenous PSD-95 (red). **F)** Quantification of the number of the total amount of PSD-95 labeled by PSD-95.FingR-tagRFP before and after expression of PSD-95.FingR-HA-RING_MDM2_ (left side). The # of puncta labeled with PSD-95.FingR-tagRFP at T0 was compared to the number of puncta immunostained for endogenous PSD-95 following expression of PSD-95.FingR-HA-RING_MDM2_ (right side). **** p < 0.0001, Mann Whitney. Error bars represent ± sem Scale bars 5 µm.

To test whether TAK243 was blocking ubiquitination, we compared western blot staining of lysates from COS7 cells expressing Ubiquitin-HA with and without TAK243 (**Fig. S1A**). Staining with anti-HA showed a distinctive laddering pattern in the lane corresponding to cells expressing Ubiquitin-HA without TAK243 consistent with ubiquitination, whereas the lanes corresponding to cells expressing Ubiquitin-HA with TAK243 and a control lane with lysate from untransfected cells showed no staining, confirming that TAK243 blocks ubiquitination. Together, our results are consistent with PSD-95.FingR-RING_Mdm2_ degrading exogenously expressed PSD-95 through ubiquitination in COS7 cells.

### Degradation of endogenous PSD-95 in neurons

To test whether PSD-95.FingR-RING_Mdm2_ can degrade endogenous neuronal PSD-95, we co-transfected doxycycline (Dox)-inducible TRE-PSD-95.FingR-HA-RING_Mdm2_ and transcriptionally regulated PSD-95.FingR-tagRFP in 14 DIV (days in vitro) cultures of rat cortical neurons. Note that PSD-95.FingR-tagRFP efficiently labels endogenous PSD-95, allowing its spatial distribution to be mapped in real-time in living cells^3^. Furthermore, transcriptional regulation matches the expression level of PSD-95.FingR with that of endogenous PSD-95, facilitating labeling with very low background. Following incubation for four days, we imaged the neurons for PSD-95.FingR-tagRFP and subsequently induced the expression of PSD-95.FingR-HA-RING_Mdm2_ with Dox. After 48 hr, we reimaged the neurons for PSD-95.FingR-tagRFP and then fixed and stained them with anti-PSD-95 and anti-HA. We found that PSD-95.FingR-tagRFP labeling was reduced by 85% (p = 0.002, Wilcoxon, n = 9 cells, 3 independent experiments (**Fig. 1C, D**) consistent with efficient ablation of endogenous PSD-95. However, when we counted the number of puncta labeled with PSD-95.FingR at T0 and compared that to the number of puncta labeled with immunostaining of endogenous PSD-95 in the same cells at the end of the experiment, it showed a reduction of only 52% (**Fig. 1E, F** p < 0.0001, Mann Whitney, n = 9 cells, 3 independent experiments). A strategy for improving this relatively low ablation rate might be provided by the results of experiments where the expression of MEF2 was found to cause the elimination of excitatory synapses^17^. In those experiments, efficient synapse elimination was found to require a combination of ubiquitination mediated by Mdm2 and interaction with the proteasome, which was mediated by Protocadherin 10 (PCDH 10). PCDH 10 is a Ca^2+^-dependent cell adhesion protein that binds to both PSD-95 and the proteasome via its proteasome interacting region (PIR)^17^. Therefore, we reasoned that adding the PIR domain to the PSD-95.FingR-RING_Mdm2_ complex might increase the efficiency with which PSD-95 is ablated.

### Optimization of degradation using the Proteasome Interacting Region

We generated a new protein, PSD-95.FingR-RING_Mdm2_-PIR, which we called PFE3. To test PFE3, we co-transfected TRE-PFE3-HA with PSD-95.FingR-tagRFP in dissociated cultures of rat cortical neurons. Following four days of incubation, we imaged the PSD-95.FingR-tagRFP and induced the expression of PFE3-HA with Dox. The neurons were imaged 48 hr after induction of TRE-PFE3 and subsequently fixed and immunostained for endogenous PSD-95 and HA (**Fig. 2A-C**). By comparing images at T0 and 48 hr, we found that the expression of PFE3 reduced the labeling of PSD-95.FingR-tagRFP by 65%, a significant difference (**Fig. 2A, B, D**, n = 14 cells, 3 distinct experiments, p < 0.0001, Mann Whitney). When we checked these results by counting PSD-95 puncta labeled with PSD-95.FingR-tagRFP at T0 and immunocytochemistry against endogenous PSD-95 at 48 hr (**Fig. 2C**), we found that Dox-inducible PFE3 reduced the number of endogenous PSD-95 puncta by 73% (**Fig. 2D**, p < 0.0001, Mann-Whitney). Thus, PFE3 consistently and efficiently reduced both total labeling of PSD-95 by the PSD-95.FingR, and the number of endogenous PSD-95 puncta. As a control, we induced RandE3 (Random.FingR-RING_Mdm2_-PIR), which contained a fibronectin scaffold with a random binding pocket instead of PSD-95.FingR^1^. The expression of RandE3 did not significantly affect PSD-95.FingR labeling (an increase of 30%, p > 0.1, Wilcoxon, n = 8, 3 independent experiments) or the number of endogenous PSD-95 puncta (an increase of 2%, p > 0.9, Wilcoxon, n = 8, 3 independent experiments, **Fig. S2A-D**).

**Figure 2:**
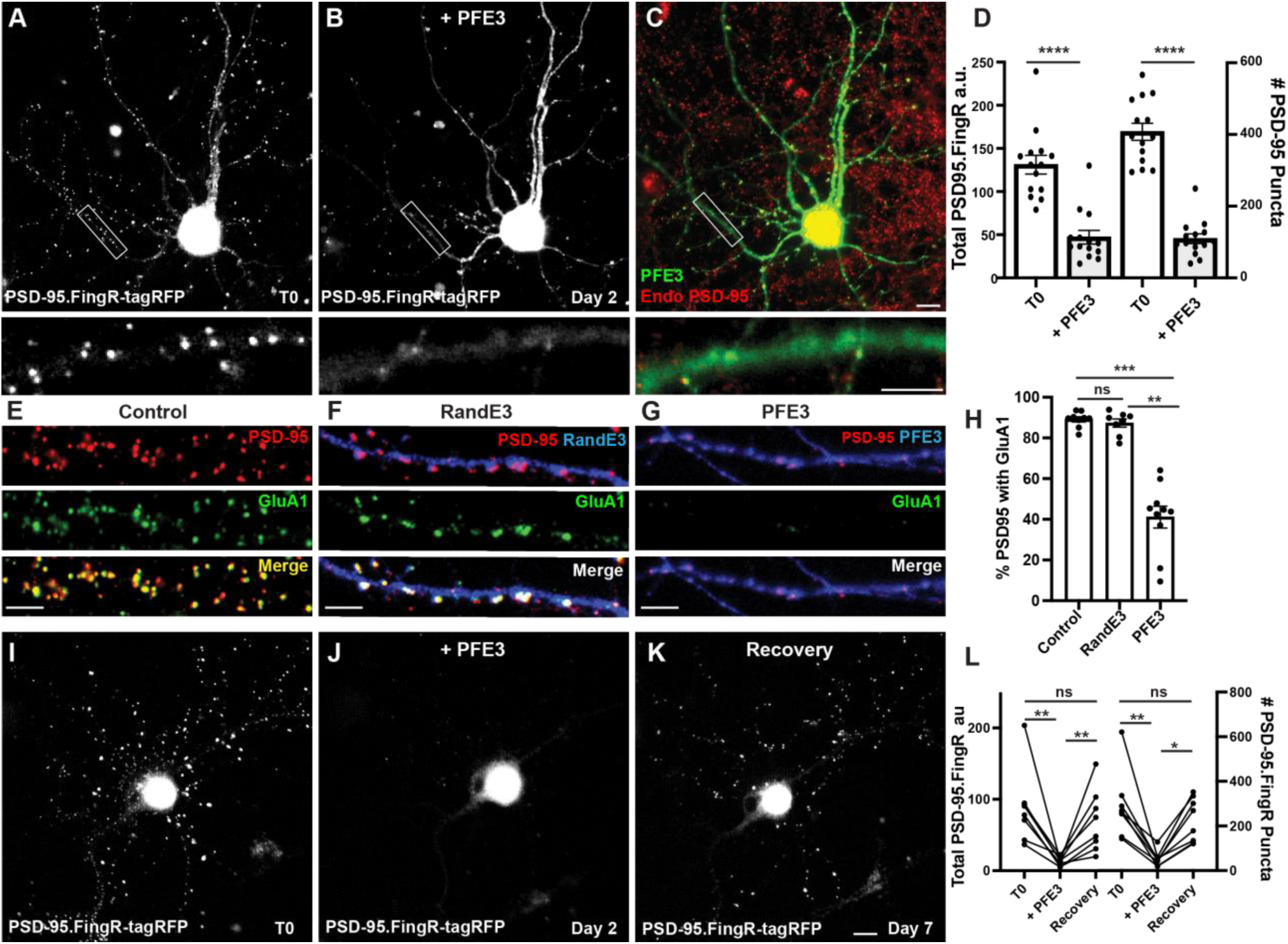
PFE3 reversibly ablates PSD-95 in neurons. **A)** Cultured cortical neuron expressing PSD-95.FingR-tagRFP before induction of PFE3. Closeup of the boxed area shown below. **B)** Same neuron as in A) after expression of PFE3 for 48 hr shows a dramatic reduction in PSD-95.FingR labeling. Closeup of the boxed area shown below confirms the lack of punctate labeling by PSD-95.FingR-tagRFP. **C)** Neuron in B) immunostained for PFE3 (green) and endogenous PSD-95 (red) confirms the lack of punctate labeling by PSD-95.FingR. Closeup of the boxed area shown below. **D)** Quantifications of the total amount of PSD-95.FingR labeling and the # of PSD-95 puncta (T0, # of puncta labeled with PSD95.FingR; + PFE3, # of puncta labeled with immunocytochemistry) show significant reductions following expression of PFE3. **** p < 0.0001, Mann Whitney. **E-G)** Cultured cortical neurons immunostained for endogenous PSD-95 in red and GluA1 in green. **E**) Untransfected. **F**) Following RandE3 (blue) expression for 48 hr, GluA1, and PSD-95 staining is intact. **G)** Following PFE3 expression (blue) for 48 hr, GluA1 and PSD-95 expression is markedly diminished. **H)** Quantification of the percentage of PSD-95 puncta positive for GluA1 staining. ns p > 0.05, *** p < 0.001, ** O < 0.01, Kruskal-Wallis multiple comparisons. **I)** Cultured neuron expressing PSD-95.FingR-tagRFP. **J)** Same neuron as in I) 48 hr after induction of PFE3 expression shows a dramatic reduction in labeling with PSD-95.FingR-tagRFP. **K)** Same neuron as in I), J) showing recovery of synapses five days after removal of PFE3. **L)** Quantification of the total amount of PSD-95.FingR-tagRFP labeling and the # of PSD-95.FingR-tagRFP-labeled puncta. ** p < 0.01, * p < 0.05, ns p > 0.05, Kruskal-Wallis multiple comparisons. Scale bars 5 µm.

To further characterize the effect of PFE3 expression, we determined how it affected the expression of AMPA receptors (AMPARs), which mediate excitatory synaptic transmission^18^. We immunostained non-transfected cultured cortical neurons for PSD-95 and GluA1, a subunit of the AMPAR, and found that 89 ± 1% of PSD-95 puncta were positive for GluA1 (**Fig. 2E, H**, 7,570 synapses, 11 cells, 3 experiments). Neurons co-transfected with the PSD-95.FingR and TRE-RandE3-HA and induced with Dox for 48 hr showed no significant change in the percentage of PSD-95-positive synapses labeled with GluA1 compared to control neurons (**Fig. 2F, H**, 87 ± 2%, 3,131 synapses, 9 cells, 2 experiments, p > 0.05, Kruskal-Wallis multiple comparisons). Quantification of GluA1 in PFE3-expressing cells showed that only 41 ± 5% (1,341 synapses, 10 cells, 2 experiments) of the remaining PSD-95 puncta also contained GluA1, a significant decrease from untransfected cells (**Fig. 2G, H,** p < 0.001, Kruskal-Wallis multiple comparisons) and RandE3 expressing cells (p < 0.01, Kruskal-Wallis multiple comparisons). Based on our result that PFE3 decreases PSD-95 puncta by 73% (**Fig. 2D**) and that only 41% of the remaining puncta are positive for GluA1, we estimate that PFE3 decreases the amount of GluA1 by approximately 90%. Thus, our experiments are consistent with PFE3 mediating a significant reduction in both PSD-95 and GluA1. Furthermore, because AMPARs are the predominant ionotropic glutamate receptor responsible for excitatory transmission^18^, our results suggest that PFE3 reduces functional excitatory connectivity.

### Degradation of PSD-95 with PFE3 is reversible

To test whether the ablation of excitatory synapses by PFE3 is reversible, we used a paradigm used previously for testing GFE3. There, we found that because GFE3 degrades itself, it is sufficient to merely cease the induction of GFE3 expression to reverse its effect^1^. Accordingly, we initially transfected TRE-PFE3-P2A-GFP and PSD-95.FingR-tagRFP and incubated for 4 days. We then induced PFE3 expression with Dox for 48 hr and then replaced the medium with Dox-free medium for an additional 5 days to allow excitatory synapses to recover. Cells were imaged immediately before adding Dox, immediately after removing Dox, and after 5 days of incubation in Dox-free medium, followed immediately by fixation and immunostaining. We found that, as in previous experiments, the induction of PFE3 reduced PSD-95.FingR-tagRFP labeling (−81 ± 5%, **Fig. 2I, J, L**, p < 0.01, Friedman multiple comparison) and a reduction in PSD-95.FingR puncta (−80 ± 5%, p < 0.01, Friedman multiple comparison, n = 8 cells, 4 experiments). Five days following removal of Dox, PSD-95.FingR labeling increased by 450 ± 100%, which was significant (**Fig. 2J-L**, p < 0.01, Friedman multiple comparison) and the number of PSD-95.FingR puncta increased by 400 ± 65% (p < 0.05, Friedman multiple comparison). PSD-95.FingR labeling was not significantly different before the addition of Dox vs. after its removal (**Fig 2L**, - 10 ± 20%, p > 0.5, Friedman multiple comparison), similarly the number of PSD-95.FingR puncta did not show a significant change before induction of PFE3 vs. after its removal (−11 ± 18%, p > 0.9, Friedman multiple comparison). Furthermore, immunocytochemistry of neurons following incubation with Dox-free media showed that endogenous PSD-95 is found on dendritic spines following synapse regrowth (**Fig. S2J**). Finally, PSD-95.FingR-tagRFP puncta showed a similar distribution overall before and after ablation (**Fig. S2K**). These results are consistent with PSD-95 degradation by PFE3 being reversible.

In a control experiment with RandE3, neurons showed no significant change in the total PSD-95.FingR labeling (+ 8 ± 6%, p > 0.9, Friedman test multiple comparisons, n = 13, 2 experiments) or the number of PSD-

95.FingR puncta (+8 ± 6%, p > 0.9, Friedman test multiple comparisons) before and after induction of RandE3 for 48 hr (**Fig. S2E-H**). Five days after the removal of Dox, both the total PSD-95.FingR labeling and the number of PSD-95.FingR puncta showed a nonsignificant decrease when compared with immediately before Dox addition (**Fig. S2F-H**, −20 ± 6%, p > 0.05, and −15 ± 5%, p > 0.2, respectively, Friedman multiple comparisons). Comparing the PSD-95.FingR before induction of RandE3 at T0 and 5 days after removal of Dox, we found nonsignificant changes in the total amount of PSD-95.FingR labeling (**Fig. S2E, G, H** −14 ± 7%, p > 0.1, Friedman multiple comparison) and the number of PSD-95.FingR puncta (−10 ± 6%, p > 0.7, Friedman multiple comparisons). Immunocytochemistry of the cells 5 days after removal of Dox confirms the labeling by PSD-95.FingR (**Fig. S2I, J**). Thus, PSD-95 degradation only occurs when the RING_Mdm2_-PIR domains are targeted by PSD-95.FingR. Finally, cotransfection of PSD-95.FingR-tagRFP and TRE-PFE3-P2A-GFP without exposure to Dox results in labeling of PSD-95.FingR-tagRFP that is not significantly different from labeling in cultures transfected with PSD-95.FingR-tagRFP alone, both in terms of total PSD-95.FingR-tagRFP labeling (p > 0.2, Mann Whitney) and number of labeled puncta (p > 0.4, Mann Whitney, Fig. S2L).

### PFE3 expression blocks excitatory synaptic transmission

To test PFE3 *in vivo*, we examined the physiological effects of expressing it in the retinas of mice. We crossed a CCK-Cre mouse, which expresses Cre in type 6 cone bipolar cells^19^, with an Ai27 mouse expressing Channelrhodopsin 2-tdTomato (ChR2-tdTomato) in Cre-expressing cells^20^ to generate mice expressing ChR2-tdTomato in type 6 cone bipolar cells. These mice were injected intravitreally with AAVs encoding either CAG-PFE3-P2A-GFP or CAG-RandE3-P2A-GFP. After 3-4 weeks, retinas were isolated and mounted in a chamber for patch clamp recording as previously described^21^. Retinas were perfused in ACSF bubbled with 95% O_2_/ 5%CO_2_. ACET (1 µM) and L-AP4 (10 µM) were added to block synaptic transmission from photoreceptors to Off and On bipolar cells, respectively, to eliminate their natural response to light. Infected cells were identified by GFP expression (**Fig. 3A**). GFP-positive On Alpha retinal ganglion cells (α-RGCs) were identified by their large cell bodies, and their identity was confirmed by dye filling. Full-field 495 nm light (0.9 mW/cm^2^) was delivered to the retina for 500 ms to trigger EPSCs (**Fig. 3A**). Synaptic currents were evoked by optogenetic stimulation of type 6 bipolar cells, which provide approximately 70% of the total synaptic input to On α-RGCs^22^. At a holding potential of −60 mV, these synaptic currents are almost purely AMPAergic, as the AMPAR antagonist DNQX blocks the response^23^. Compared to RGCs infected with the control virus, RGCs infected with the PFE3 virus had EPSCs with > 80% reduction in amplitude, 79 ± 30 pA (n = 9 cells, 5 mice) vs. 445 ± 78 pA (n = 8 cells, 3 mice), a significant difference (p < 0.001, Mann Whitney, **Fig. 3B-D**), which is consistent with a reduction in postsynaptic receptor density. Overall, our results are consistent with PFE3 expression leading to the degradation of PSD-95, which, in turn, reduces the number of AMPA receptors at excitatory synapses, causing a reduction in synaptic transmission.

**Fig. 3.**
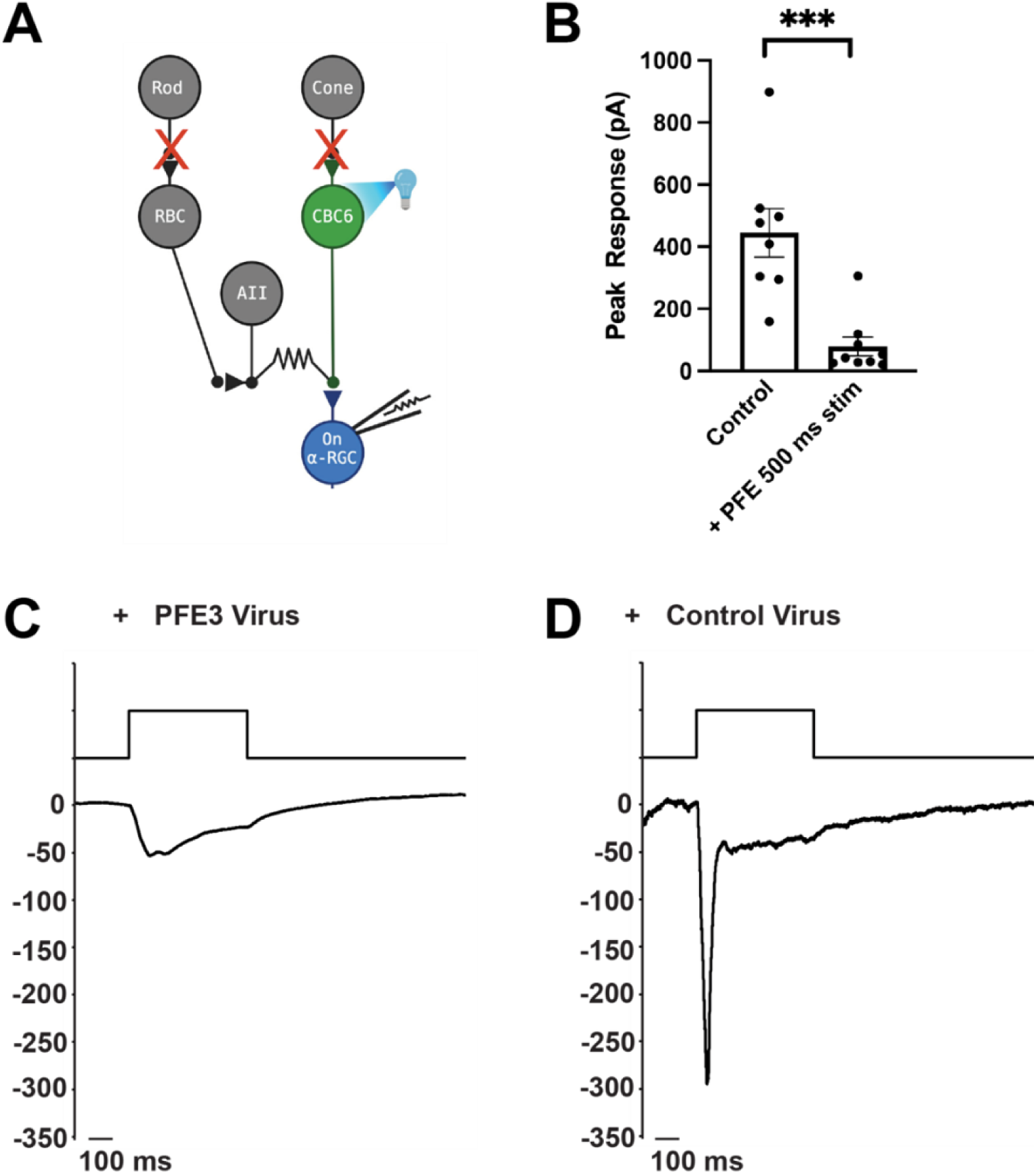
Expression of PFE3 reduces synaptic transmission in vivo. **A)** Schematic depiction of the retinal circuit used to assess AMPA receptor function. Channelrhodopsin-2 is expressed in type 6 cone bipolar cells (CBC6), which are presynaptic to the On α-retinal ganglion cell (α-RGC). Synaptic output of photoreceptors is pharmacologically blocked. **B)** Population data comparing peak EPSC amplitude for retinas from mice infected with PFE3 and control viruses. Responses were generated with a 495 nm, 500 ms flash of light. Symbols are values for individual cells. *** p < 0.001, Mann Whitney. Error bars represent ± s.e.m. **C)-D)** Sample whole cell patch clamp recording from an On α-RGC infected with the PFE3 virus (C) or a RandE3 control virus (D).

### Photoactivatable degradation of inhibitory synapses

To generate a photoactivatable version of GFE3 to ablate inhibitory synapses, we first used existing photodimerization systems, such as the Cryptochrome (Cry2) system, but we found them to have substantial background activity leading to loss of inhibitory synapses even in the dark (**Fig. S4A**). To overcome this deficiency, we developed a new photoactivatable complex based on the intrinsically photocleavable protein PhoCl. PhoCl undergoes cleavage when exposed to violet (∼400 nm) light, creating two peptides, the C-terminal ten amino acids, including the chromophore, and the remainder of the protein, referred to as the “empty barrel”^13^. Following photocleavage, a novel epitope is exposed on the empty barrel. Based on this property of PhoCl, we developed a photoactivatable complex, PhLIC (**Ph**oCl-based **L**ight-**I**nducible **C**omplex), consisting of the empty barrel and a 10-mer peptide called PhoCl binding peptide (PBP) that binds to the empty barrel but not to full-length PhoCl (**Fig. 4A**). Because PhoCl does not undergo cleavage in the dark or ambient light^13^, there should be no background formation of PhLIC. To generate PBP, we used the *in vitro* directed evolution technique, mRNA display, which we have used to create binders to synaptic proteins^3,24,25^ (**Fig. S3A**). We first constructed a library of ∼10^14^ semi-random 10-mers, where each amino acid was biased towards the corresponding amino acid in the original C-terminus of PhoCl. Following the first selection, which lasted six rounds, we found PBPs that bound to cleaved PhoCl in COS7 cells but not in neurons (data not shown). We then remade the library by amplifying the remaining PBPs after the sixth selection round and performing another selection.

**Figure 4:**
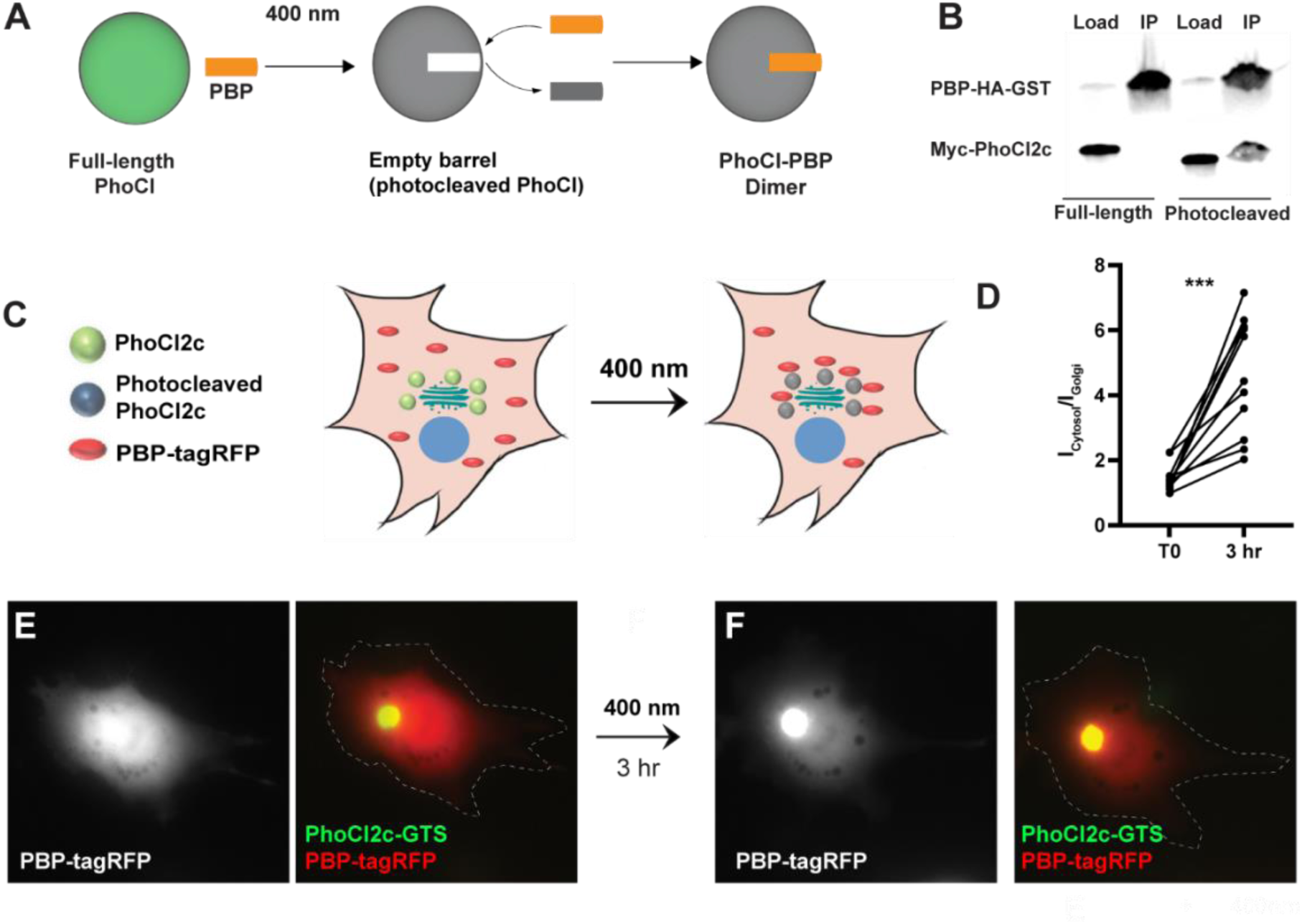
Testing of PhLIC, a photoactivatable dimer. **A)** Schematic of PhLIC (**Ph**oCl-based **L**ight-**I**nducible **C**omplex). Full-length PhoCl is photocleaved with 400 nm light, exposing an epitope recognized by the PhoCl Binding Peptide (PBP), leading to dimerization of photocleaved PhoCl and PBP. **B)** COS7 lysate expressing PBP-HA-GST was incubated with purified full-length or photocleaved Myc-PhoCl2c. PBP-HA-GST pulls down photocleaved but not full-length PhoCl2c. **C)** Schematic of Golgi-targeting assay. COS7 cell expressing Golgi-targeted PhoCl2c in green and PBP-tagRFP in red before (left) and 3 hr after illumination with 400 nm light for 10 sec every 30 sec for 3 min (right). **D)** Quantification of the ratio of the total fluorescence associated with PBP-tagRFP localized in the cytosol (I_cytosol_) vs. at the Golgi (I_Golgi_) before and 3 hr after photocleaving PhoCl2c. *** p < 0.001, Wilcoxon. **E)** COS7 cell before photoactivation showing PBP-tagRFP (left) and a merge (right) of PBP-tagRFP (red) and PhoCl2c-GTS (green). **F)** Same COS7 cell as in E) following photoactivation with 400 nm light for 1 min showing PBP-tagRFP (left) and a merge (right) of PBP-tagRFP (red) and PhoCl2c (green). Scale bar represents 5 µm.

We tested the results of this second selection by generating full-length and photocleaved PhoCl2c (see methods) and then pulling it down with PBP. Note that PhoCl2c is an improved version of the original PhoCl containing 5 mutations and with faster and more efficient cleavage^12^. We found that PBP pulled down photocleaved PhoCl2c, but not full-length PhoCl2c (**Fig. 4B**), suggesting that PhoCl2c/PBP complex, which we called PhLIC (**Ph**oCl-based **L**ight-**I**nducible **C**omplex), could act as a photoactivatable system with low background. We further tested PhLIC by co-transfecting COS7 cells with Golgi-targeted PhoCl2c (GTS-PhoCl2c) and PBP-tagRFP and exposing them to 400 nm light. If PBP binds to photocleaved PhoCl2c but not full-length PhoCl2c, we would expect PBP-tagRFP to be expressed diffusely throughout the cell initially and then translocate to the Golgi after photocleavage of PhoCl2c (**Fig. 4C**). Following expression in COS-7 cells, GTS-PhoCl2c localized in a concentrated oval in the center of the cell consistent with Golgi localization. In contrast, PBP-tagRFP localized diffusely throughout the cytoplasm, with a ratio of labeling in the area colocalized with GTS-PhoCl2c vs. the rest of the cell of 1.5 ± 0.1 (n = 11 cells, 2 experiments, **Fig. 4D, E**). Following exposure to 400 nm light for 1 min and incubation in the dark for 3 hr, PBP-tagRFP became 3.3 ± 0.5 times more concentrated in the area colocalized with GTS-PhoCl2c staining than vs. the rest of the cell, a significant difference (P < 0.001, n = 11, Wilcoxon, 2 experiments **Fig. 4D, F**). Note that exposure to ambient light did not affect the localization of PBP-tagRFP (**Fig. S3B-E**), and exposing cells transfected only with tagRFP to 400 nm did not affect the localization of tagRFP (**Fig. S3F-I**). These experiments suggest that PhLIC shows robust activation upon exposure to 400 nm light but no activation in ambient light.

### Photoactivatable GFE3

To generate a photoactivatable version of GFE3 using PhLIC (paGFE3), we fused PhoCl2c to transcriptionally controlled Gephyrin FingR (tagRFP-GPHN.FingR-PhoCl2c) and PBP to RING_XIAP_ (PBP-E3). Note that tagRFP-GPHN.FingR-PhoCl2c performs two functions. It is targeted to inhibitory synapses, where it acts as an anchor that recruits PBP-E3 once PhoCl2c has been cleaved. It also acts as a label, allowing inhibitory synapses to be visualized before and after PhCl2c is cleaved (**Fig. 5A**). To test whether paGFE3 labels inhibitory synapses, we expressed tagRFP-GPHN.FingR-PhoCl in cortical neurons in culture and co-labeled tagRFP and endogenous Gephyrin. The two labels are colocalized, consistent with tagRFP-GPHN.FingR-PhoCl labeling inhibitory synapses (**Fig. S4B**). To test if paGFE3 caused background degradation of inhibitory synapses, we compared cultured cortical neurons transfected with paGFE3 (transcriptionally regulated tagRFP-GPHN.FingR-PhoCl and PBP-E3) with similar neurons transfected with tagRFP-GPHN.FingR-PhoCl alone and incubated for 5 days. Quantification of tagRFP labeling showed an 8% reduction, which was not significant (n = 10 cells, 1 experiment, p > 0.4, Mann Whitney, **Fig. S4A**), consistent with no background degradation of Gephyrin in the dark. In contrast, when Cry2-tagRFP-GPHN.FingR and CIB1-RING_XIAP_ were co-transfected into cultured cortical neurons and incubated for five days, tagRFP labeling was 48% less than when GPHN.FingR-Cry2-tagRFP was transfected by itself (650 ± 90, au, n = 10 cells, 1 experiment vs.1250 ± 110 au, n = 9 cells, 1 experiment), a significant difference (p < 0.001, Mann Whitney, **Fig. S4A**).

**Fig 5.**
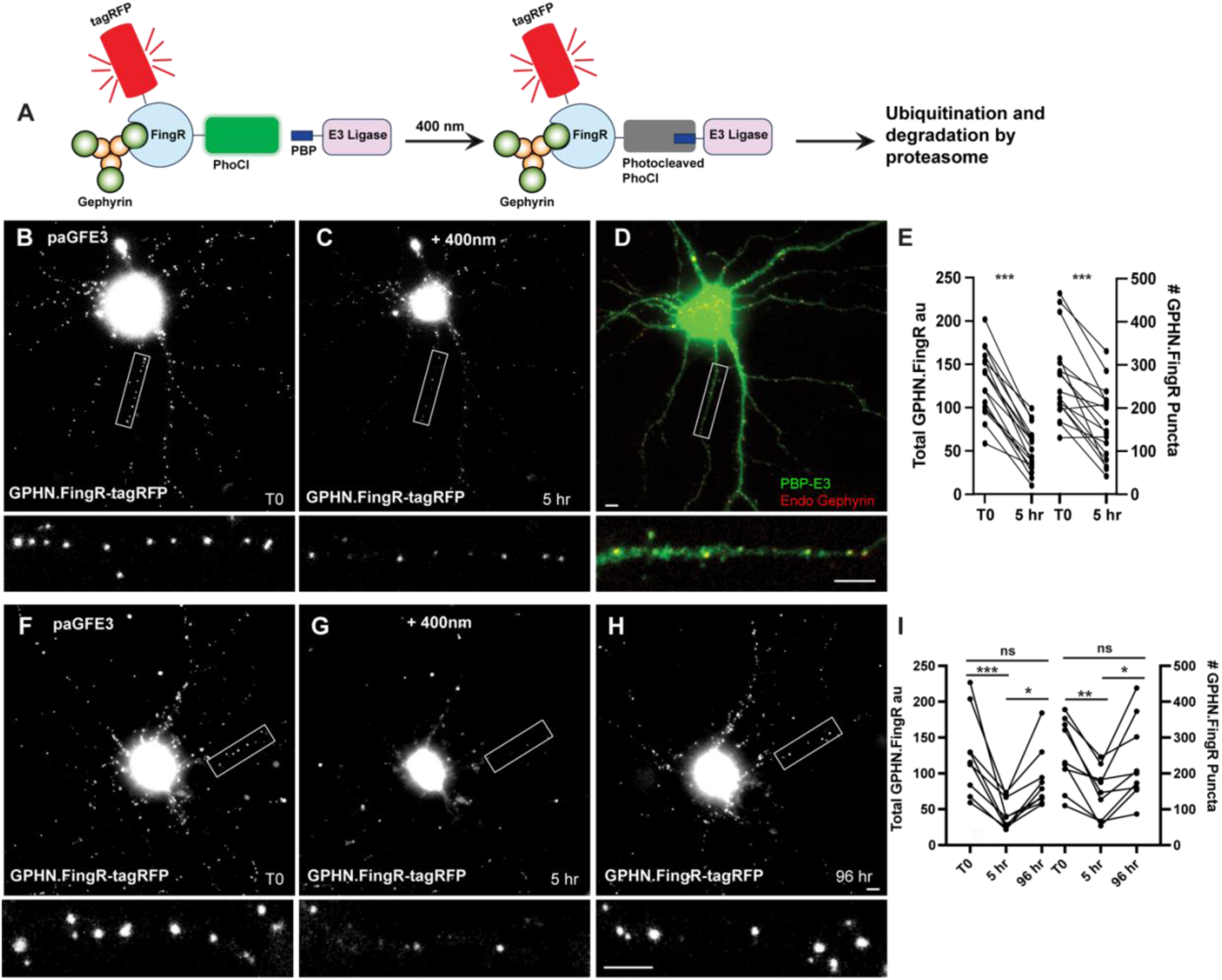
Reversible optogenetic ablation of inhibitory synapses with pa-GFE3. **A)** Schematic of paGFE3. tagRFP-GPHN.FingR-PhoCl2c binds to Gephyrin. After photocleavage with 400 nm light, PBP binds photocleaved PhoCl, recruiting the E3 ligase domain, which ubiquitinates Gephyrin and targets it for degradation by the proteasome. **B)** Cultured cortical neuron expressing tagRFP-GPHN.FingR-PhoCl2c + PBP-HA-E3 (pa-GFE3) for 5 days. Closeup of the boxed area shown below. **C)** Same neuron as in B) 5 hr after illumination with 400 nm light for 1 min showing reduced labeling with tagRFP-GPHN.FingR-PhoCl2c. Closeup of the boxed area shown below. **D)** Neuron in C) immunostained for PBP-HA-E3 (green) and endogenous Gephyrin (red) showing sparse labeling for endogenous Gephyrin. Closeup of the boxed area shown below. **E)** Quantification of the total amount of GPHN.FingR and number of GPHN.FingR puncta before and 5 hr after illumination with 400 nm light. *** p < 0. 001, Wilcoxon. **F)** Cultured cortical neuron expressing pa-GFE3. Closeup of the boxed area shown below. **G)** Same neuron as in F) 5 hr after illumination with 400 nm light for 1 min showing reduced labeling with tagRFP-GPHN.FingR-PhoCl2c. Closeup of the boxed area shown below. **H)** Same neuron as in F), G) 4 days after illumination showing a recovery of tagRFP-GPHN.FingR-PhoCl2c labeling of synapses. Closeup of the boxed area shown below. **I)** Quantification of the total amount of tagRFP-GPHN.FingR-PhoCl2c labeling and number of GPHN.FingR puncta showing recovery of synapses. *** p < 0.001, ** p < 0.01, * p < 0.05, ns p > 0.1, Friedman test multiple comparisons. Scale bars represent 5 µm.

To test whether the paGFE3 could mediate light-dependent degradation of Gephyrin, we expressed both components of paGFE3 (tagRFP-GPHN.FingR-PhoCl2c and PBP-E3) in cultured cortical neurons and incubated them for five days. Subsequently, neurons were intermittently exposed to 400 nm light for 1 min (see methods). After 5 hr, the cultures were reimaged. The neurons showed a 61 ± 3% reduction in tagRFP labeling (130 ± 9 a.u. vs. 52 ± 6 a.u., a significant difference, p < 0.001, Wilcoxon, n=17 cells, 3 experiments, **Fig. 5B-E**) and a 40 ± 6% drop in the number of GPHN.FingR puncta (268 ± 24 vs. 161 ± 20 puncta), a significant change (p < 0.001, Wilcoxon, n=17, 3 experiments, **Fig. 5B-E**). Importantly, cells expressing transcriptionally regulated tagRFP-GPHN.FingR-PhoCl2c without PBP-RING had a much smaller reduction in labeling (10 ± 2%, **Fig. S4C-E**) when exposed to 400 nm light as above, although it was statistically significant (p < 0.001, Wilcoxon, n=14 cells, 3 experiments, **Fig. S4**). However, the number of Gephyrin puncta labeled by the GPHN.FingR did not show a significant change (0.4 ± 2%, p > 0.7, Wilcoxon, n = 14 cells, 3 experiments, **Fig. S4E**), indicating the reduction in total GPHN.FingR labeling is likely a result of photobleaching and not due to the decrease in Gephyrin. Thus, our results are consistent with cells expressing paGFE3 undergoing efficient degradation of Gephyrin puncta when exposed to 400 nm light.

To determine whether Gephyrin ablation mediated by paGFE3 is reversible, we transfected both components of pa-GFE3 in dissociated cortical neurons as before. Four days after transfection, cells were imaged and then exposed to 400 nm light for 1 minute at time point T0. Five hr later, the cells were reimaged and incubated for another four days, at which point they were imaged again. We reasoned that the PhLIC-GFE3 complexes would be self-degrading, as was GFE3. Thus, in the absence of light, they should be degraded and allow regeneration of Gephyrin puncta after a sufficient waiting period. Comparing the images, we found the total fluorescence from the tagRFP-GPHN.FingR-PhoCl2c label was reduced by 63 ± 5% between T0 and +5 hr, then increased by 130 ± 28% between +5 hr and +4 day, a significant difference (p < 0.001 for T0 to +5 hr; p < 0.05, +5 hr to +4 days, Friedman test, multiple comparisons, n = 9 cells, 2 experiments, **Fig. 5F-I**). In addition, there was a 20 ± 9% decrease from T0 to +4 days, which was not significant (p > 0.1, Friedman test, multiple comparisons, **Fig. 5I**), suggesting that paGFE3 is reversible. We found similar results when we counted the number of GPHN.FingR puncta which showed a 43 ± 7% reduction (p < 0.01, Friedman test, multiple comparisons) followed by a 76 ± 22% increase (P < 0.05, Friedman test, multiple comparisons), representing a nonsignificant 6 ± 12% decrease from the original (p > 0.99, Friedman test, multiple comparisons, n = 9 cells, 2 experiments). We also found that puncta before and after ablation were in similar locations (**Fig. S4F**).Together, our results indicate that paGFE3 can mediate light-inducible degradation of endogenous Gephyrin that has virtually no background and is reversible.

### Chemically induced GFE3

We also generated a chemogenetic version of GFE3 (chGFE3) that can be used when an inducible version of GFE3 is needed, but paGFE3 is unsuitable. We used a chemically induced dimerization system based on E. coli dihydrofolate reductase (eDHFR) and the HaloTag protein. When combined with the small molecule TH, a fusion of trimethoprim (TMP) and HaloTag ligand, eDHFR and HaloTag form a bio-orthogonal complex in eukaryotic cells^27^. Furthermore, the formation of the complex is reversible because the binding of eDHFR to TMP is not covalent and can be outcompeted by adding excess TMP. We divided GFE3 into its two principal components, GPHN.FingR and RING_XIAP_ and fused transcriptionally regulated GPHN.FingR to the HaloTag peptide and RING_XIAP_ to eDHFR, to give tagRFP-GPHN.FingR-HaloTag and eDHFR-E3 (**Fig. 6A**). As with tagRFP-GPHN.FingR-PhoCl2c, tagRFP-GPHN.FingR-HaloTag will both label inhibitory synapses and provide an anchor to which E3 can be recruited to initiate the degradation of Gephyrin. To test whether recruiting eDHFR-E3 to inhibitory synapses would lead to the loss of Gephyrin, we transfected cultured neurons with transcriptionally controlled tagRFP-GPHN.FingR-HaloTag and DHFR-E3 (chGFE3). After visualizing Gephyrin puncta, neurons were treated with 100 nM TH to induce dimerization. Incubation of the neurons with TH for 4 hr showed a reduction of Gephyrin by 73% ± 3% (p < 0.001, Wilcoxon, **Fig. 6B-E** n = 12 cells, 3 experiments) and a reduction in the number of GPHN.FingR puncta by 53 ± 7% (p < 0.001, Wilcoxon, n = 12 cells, 3 experiments). Incubating the neurons with TH for 24 hr reduced total Gephryin labeling by 89 ± 2%, a significant reduction (p < 0.001, Wilcoxon, n = 12 cells, 3 experiments **Fig. 6F-I**) and the number of GPHN.FingR puncta by 92 ± 0.1% (p < 0.001, Wilcoxon, n = 12 cells, 3 experiments). Immunocytochemistry against endogenous Gephyrin confirms the loss of inhibitory synapses at 4 hr and 24 hr (**Fig. 6D, H**). To confirm that Gephyrin loss depends on the dimerization between the GPHN.FingR and RING_XIAP_ neurons were transfected with tagRFP-GPHN.FingR-HaloTag and SnapTag-E3 and treated with TH. There were nonsignificant increases in labeling with GPHN.FingR (+ 20 ± 16%, p > 0.5, Wilcoxon, n = 8, 2 experiments), and the number of GPHN.FingR puncta (+ 11 ± 10%, p > 0.99, Wilcoxon, n = 9, 2 experiments, **Fig.S5A-C)**. Endogenous Gephyrin was also unaffected in these cells **(Fig. S5D)**. Thus, our results are consistent with chGFE3 mediating efficient chemically-induced degradation of Gephyrin.

**Figure 6:**
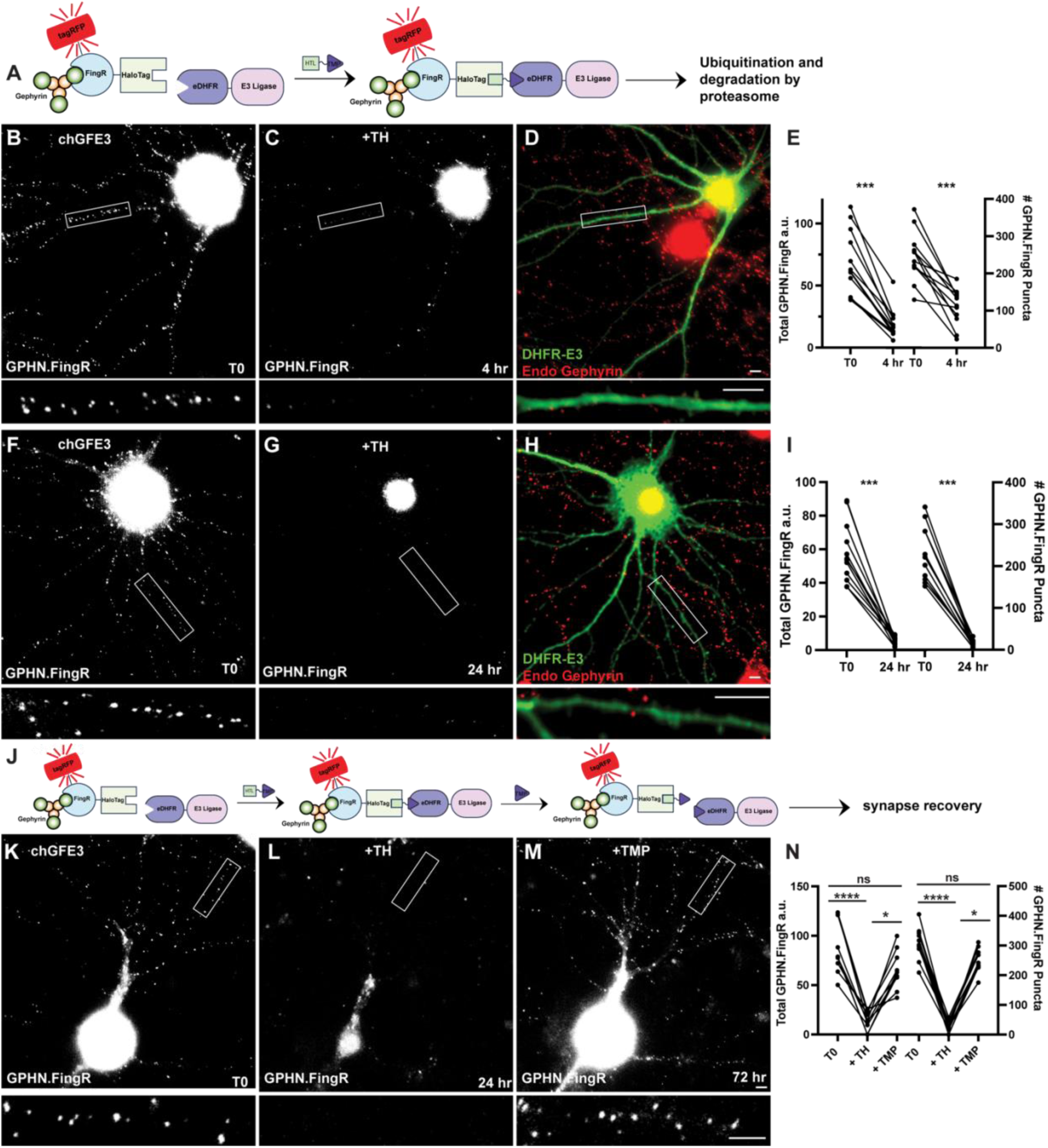
chGFE3 reversibly ablates Gephyrin. **A)** Schematic of chGFE3. The addition of TMP-HaloTag ligand (TH) dimerizes HaloTag and eDHFR, leading to the recruitment of the E3 ligase to Gephyrin, which ubiquitinates and degrades it. **B)** Cultured cortical neuron expressing GPHN.FingR-HaloTag and eDHFR-RING_XIAP_ (chGFE3) for 4 days. Closeup of the boxed area shown below. **C)** Same neuron as in B) 4 hr after addition of 100 nM TH. Closeup of the boxed area shown below. **D)** Immunocytochemistry of the neuron in C) for endogenous Gephyrin (red) and eDHFR-E3_XIAP_ (green). Note the lack of red puncta on dendrites labeled green. Closeup of the boxed area shown below. **E)** Quantification of total Gephyrin labeled by GPHN.FingR-tagRFP and number of GPHN.FingR puncta after 4 hr of incubation with 100 nM TH. *** p < 0.001, Wilcoxon. **F)** Cultured cortical neuron expressing GPHN.FingR-HaloTag and eDHFR-RING for 4 days. Closeup of the boxed area shown below. **G)** Same neuron as in F) after 24 hr incubation with 100 nm TH showing loss of Gephyrin. Closeup of the boxed area shown below. **H)** Neuron in G) immunostained for endogenous Gephyrin (red) and DHFR-E3 (green). Closeup of the boxed area shown below. **I)** Quantification of the total amount of Gephyrin labeling by the GPHN.FingR after the addition of TH. *** p < 0.0001, Friedman test multiple comparisons. **J)** Schematic illustrating dissociation of chGFE3 with the addition of TMP. **K)** Cultured cortical neuron expressing GPHN.FingR-HaloTag and eDHFR-RING_XIAP_ for 4 days. Closeup of the boxed area shown below. **L)** Same neuron as in K) after 24 hr incubation with 100 nm TH showing loss of Gephyrin. Closeup of the boxed area shown below. **M)** Same neuron as in L) after 48 hr incubation with 100 µM TMP showing recovery of Gephyrin puncta. Closeup of the boxed area shown below. **N)** Quantification of the total amount of Gephyrin labeling by the GPHN.FingR after the addition of TH and then TMP. **** p < 0.0001, * p < 0.05, ns p > 0.2 Friedman Test multiple comparisons. Scale bar represents 5 μm.

The tagRFP-GPHN.FingR-Halotag/eDHFR-E3/TH complex should dissociate with the addition of excess TMP and reverse chGFE3 (**Fig. 6J**). To test whether chGFE3 is reversible, we transfected cultured cortical neurons with GPHN.FingR-HaloTag and eDHFR-RING and treated with TH for 24 hr to induce degradation of Gephyrin, which led to a reduction of 83 ± 4% in GPHN.FingR-HaloTag labeling and 91 ± 2% reduction in the number of GPHN.FingR puncta (**Fig. 6K, L, N,** p < 0.0001 for both, Friedman test, multiple comparisons, n = 10 cells, 2 experiments). We then added 100 µM TMP to the media for 24 hr to compete with TH to dissociate GPHN.FingR-RING_XIAP_ dimers and block the formation of new dimers (**Fig. 6M**). Imaging the neurons 48 hr after adding TMP later showed a 1450 ± 760% increase in the total GPHN.FingR labeling and 1168 ± 200% increase in the number of Gephyrin puncta at 72 hr, both of which were significant (p < 0.05 for both, Friedman multiple comparisons, **Fig. 6M, N)**. The amount of GPHN.FingR labeling at 72 hr compared to labeling at T0 was reduced by 24 ± 5%, and the number of GPHN.FingR puncta decreased by 21 ± 4%, both of which were not significant (p > 0.2, Friedman multiple comparison, **Fig. 6K, M, N**). Furthermore, immunocytochemistry shows colocalization between the GPHN.FingR and the GABA_A_ receptor, the main ionotropic GABA receptor^28^, denoting a functional synapse **(Fig. S5E)**. Finally, we found that the Gephyrin puncta before and after ablation had similar distributions across dendrites (**Fig. S5F**). Thus, our results suggest that in neurons expressing chGFE3, adding TH causes ablation of inhibitory neurons, and adding TMP reverses the effect.

## Discussion

In this study, we developed three new synapse ablators: PFE3, which ablates excitatory synapses; paGFE3, a photoactivatable version of the inhibitory synapse ablator; GFE3; and chGFE3, a chemically activated version of GFE3. All three tools ablate specific postsynaptic sites and can be expressed in particular cell types, allowing circuit breaking with two-fold specificity.

PFE3 is composed of three distinct functional domains. Ubiquitination of PSD-95 is mediated by the E3 ligase Mdm2, the PIR domain recruits proteasomes to the synapse to degrade the ubiquitinated protein, and both domains are targeted to PSD-95 through PSD-95.FingR. Its design is based on work showing that coordinated actions of Mdm2 and PIR from Protocadherin 10 mediate the degradation of excitatory synapses^17^. These findings were corroborated when we found efficient degradation of PSD-95 and ablation of synapses was only achieved if all three components were included in PFE3. Expression of PFE3 caused highly efficient ablation of PSD-95 in cultured cortical neurons and blocked excitatory synaptic transmission in retinal ganglion cells in vivo.

Efficient elimination of excitatory synapses was achieved after 48 hr of PFE3 expression, which contrasts with PSD-95 knockout mice, where reductions in AMPA receptors and excitatory transmission were either not significant or small depending on the stage of development. One reason for this discrepancy could be that transcription of the MAGUK protein SAP-102, which is highly homologous to PSD-95, is upregulated in PSD-95 knockout mice and could provide compensation^29^. PSD-95.FingR labels SAP-102^3^, which suggests that PFE3 would degrade any SAP-102 that might otherwise compensate for the reduction in PSD-95. Thus, in the context of PFE3, the binding of PSD-95.FingR to MAGUKs homologous to PSD-95, such as SAP-102 and SAP97, could enable more effective and widespread ablation of excitatory synapses than if it bound only to PSD-95. Alternatively, compensatory changes in transcription could be relatively slow, or they respond specifically to changes in mRNA levels rather than protein levels. In the future, it should be possible to use PFE3 to explore mechanisms of compensation that maintain homeostasis when protein levels change.

Another obvious application of PFE3 is to probe the role of PSD-95 in maintaining synaptic structure. For instance, what is the relationship between PSD-95 and dendritic spines? It would also be interesting to closely examine how synapses change before and after ablation of PSD-95. PSD-95 puncta grow back in cultured neurons, but what would happen in mature neurons in vivo? If they grow back, would they reconnect with their original presynaptic partners, and if so, would neural circuit function be changed following regrowth? It would also be interesting to see whether it is possible to permanently alter the connectivity of circuits after extended periods of expression of PFE3.

We added temporal and spatial control capability to GFE3 by enabling its activation with light (paGFE3) or chemicals (chGFE3). We used a standard method for regulating a protein where two distinct domains that are necessary and sufficient for protein function are fused to separate components of a light- or chemically-induced protein complex. Previously, this approach has been used for controlling transcription^26^ neurotransmitter release^7^, gene editing^30^, recombination^31^, and protease cleavage^32^. However, background activation in photoactivatable complexes based on photoreceptors, such as cryptochrome 2^26^, phototropin^33^, or Vivid^34^, makes them unsuitable for regulating GFE3, which is highly potent. As an alternative, we developed a novel photoactivatable complex, PhLIC, based on the photocleavable protein PhoCl and the 10-mer PBP, which exhibited no measurable background when tested in the context of paGFE3.

This lack of background would make paGFE3 ideal for use in transgenic animals such as mice or zebrafish, where the complex would have to remain inert for long periods before activation. Expressing paGFE3 in a Gal4 (zebrafish) or Cre (mouse) driver line would make it possible to ablate inhibitory synapses in specific cell types with precise temporal control and without injecting viruses. In addition, with precise application of 400 nm light, it could be possible to ablate inhibitory synapses with single-cell spatial resolution. Furthermore, the PhLIC complex could be used to make photoactivatable versions of proteins other than GFE3. For instance, it could be used to create a split version of a potent toxin, which requires that no background activity be present until photoactivation causes it to kill or alter the function of specific cells at a particular time.

chGFE3 is based on a system similar to one for degrading exogenous proteins with eDHFR tags^35^. Our system has the same benefits of efficiency and reversibility, but it can target endogenous Gephyrin and ablate inhibitory synapses. In addition, we showed that, like paGFE3, chGFE3 has no background. Although its desirable qualities overlap with those of paGFE3, it could be used in contexts where it would be too difficult or time-consuming to photoactivate GFE3. For instance, if the target cells were spread over a relatively large region or when sustained ablation is necessary. One potential drawback to applying chGFE3 in vivo is the necessity of injecting the chemical dimerizer TH. However, it may be that TH can be administered systemically, particularly since both components of TH (the HaloTag ligand and TMP) can each penetrate the brain when injected systemically^36,37^.

In conclusion, we have generated novel proteins for ablating excitatory and inhibitory synapses, which can be used to probe the structure and function of neural circuits.

## Acknowledgments

We thank all Arnold lab members for helpful discussions. We also thank Alexandra Delgadillo, Jacqueline Rivera, and Ben Shapero for technical assistance. We thank Scott Nawy from the Kramer lab for compiling the data in Fig. 3. This work was supported by a grant (NS115610) to D.A. from NINDS and the Brain Initiative. mRNA display was originally developed by Richard Roberts, who helped establish it in the Arnold lab.

## Author Contributions

D.A. conceived the study. D.A. and A.B. designed the experiments and wrote the paper. A.B. performed all experiments except for those involving electrophysiology. A.B. and S.D. analyzed the data. R.K. and S.Y. designed the electrophysiology experiments, and S.Y. performed them. C.D. and D.C. provided reagents for chGFE3 and helped with the design of chGFE3-related experiments. R.E.C. developed PhoCl and PhoCl2c and R.E.C., X.C., and W.E. provided cleaved PhoCl protein for the mRNA display selection of PBP.

## Methods

### Preparation of cultured cortical neurons

Experimental protocols were conducted according to the National Institutes of Health guidelines for animal research and were approved by the Institutional Animal Care and Use Committee at the University of Southern California.

E19 embryos were removed, and cortices were dissected in Hank’s balanced salt solution (HBSS, Thermo Fisher Scientific, cat. # 14025076) supplemented with 0.1 mM HEPES (Thermo Fisher Scientific, cat. # 15630-080). Cortices were digested with neuronal isolation enzyme with Papain (Thermo Fisher Scientific, cat # 88285) for 30 min at 37°C. The tissue was centrifuged at 1000 rpm for 1 min and washed three times with HBSS-HEPES. Subsequently, the tissue was triturated to dissociate the cells, and undissociated cells were separated out with a cell strainer (Falcon, cat # 352340). Neurons were plated at a density of 8 x 10^4^ on 2-well chambered cover glass (Cellvis, cat # C2-1.5H-N) pre-treated overnight with 0.2 mg/mL Poly-D-Lysine (Sigma-Aldrich, cat# P0899) and washed three times with water. Cells were plated in Neurobasal media (NBM, Thermo Fisher Scientific, cat # 21103-049) supplemented with 5 mL/L Glutamax (Thermo Fisher, Fisher Scientific, cat. # 35050-061), 1 mg/L gentamicin solution (Thermo Fisher Scientific, cat. # 15750-078) 10 mL/L B27 (Thermo Fisher Scientific, cat # 17504044), and 5% FBS (Hyclone, cat # SH30071.03). Cells were cultured in a 5% CO_2_ incubator at 37°C. Four hr after plating, the medium was diluted 1:3 with serum-free supplemented NBM. At 6 days *in vitro* (DIV), the medium was diluted 1:3 with supplemented NBM.

### Transfection, image capture and analysis of cultured neurons

14-15 DIV cultured cortical neurons were transfected with CalPhos mammalian transfection kit (Takara, cat # 631312). Crystals were washed and replaced with 2:1 conditioned media: fresh supplemented neurobasal media.

Live imaging of neurons was performed in Imaging Buffer consisting of HBSS (Thermo Fisher Scientific, cat # 14025076) supplemented with 0.1 mM HEPES (Thermo Fisher Scientific, cat. # 15630-080). Imaging of immunostained and live cells was done on an Olympus IX81 inverted microscope with a 60X water objective at 1.0 x zoom, an EM-CCD digital camera (Hamamatsu, cat. # C9100-02), GFP–mCherry and Cy5.5 filter cubes (Chroma Technology, cat. # 49000, cat. # 59022 and # SP105), an MS-2000 XYZ automated stage (Applied Scientific Instrumentation), an X-cite exacte mercury lamp (Excelitas Technologies) and Metamorph software (Molecular Devices).

Image analyses were performed using ImageJ software. SynQuant^38^ was used to detect PSD-95.FingR puncta for excitatory synapses and GPHN.FingR puncta for inhibitory synapses. Synapse detection by SynQuant was reviewed, and false positive signals were manually removed. The total amount of PSD-95 or Gephyrin FingR was calculated by taking the sum of the product of the area and intensity for each punctum.

### Immunocytochemistry of cultured neurons

Cells were fixed with 4% PFA (Electron Microscopy Sciences, cat. # 15714) for 5 min and washed three times for 5 min with PBS. Cells were then permeabilized and blocked with blocking buffer (1% bovine serum albumin, 5% normal goat serum, and 0.1% Triton X-100 in PBS) for 30 min. The primary antibody was then diluted in blocking buffer and added to the cells for 1 hr. After three 5 min washes with PBS, the secondary antibody was diluted in blocking buffer and added to cells for 1 hr in the dark. The cells were again washed three times with PBS and imaged. Primary antibody concentrations used were: mouse anti-PSD-95 (1:3000, Novus Biological, cat. # NB300-556), rabbit anti-GluA1 (1:1000, Millipore Sigma, ABN241), and chicken anti-GFP (1:10,000, Aves Labs, cat. # NC9510598), Rabbit anti HA (1:1000, Cell Signaling, cat. # 3724), chicken anti myc (1:1000, Novus Biologicals cat. # NB600-334). Secondary antibodies used were the following from Thermo Fisher Scientific at 1:1000: goat anti chicken-Alexa Fluor 488 (cat. # A-11039), goat anti-rabbit Alexa Fluor 594 (cat. # A-11012), goat anti-rabbit Alexa Fluor 647 (cat. # A-21245), and goat anti-mouse Alexa Fluor 594 (cat. # A-11032).

### mRNA display

mRNA display was carried out as described in ^3^ with the following modifications. The mRNA display library was constructed with a 5’ constant region including a T7 promoter, transcription start sequence and a ΔTMV translation enhancer region followed by a semi-random variable region encoding PBP followed by a 3’ constant region containing a short flexible linker, HA tag and the splint sequence used to ligate the transcript to puromycin (Keck oligonucleotide facility, Yale). The cDNA library was purified with UREA-PAGE, and Klenow extension (NEB, cat. # 0210S) was used to generate dsDNA library. The library was then transcribed with T7 RNA Polymerase and ligated to puromycin (pF30P, Oligo Synthesis at Yale School of Medicine) using T4 DNA Ligase (NEB). The mRNA-puromycin fusion was purified with Urea-PAGE and electroeluted with an Elutrap (Schleicher & Schuell BioScience). The library was translated with rabbit reticulocyte lysate (Promega cat. # L4960) and purified with dT-25 BIOTEG beads (NEB cat. # S1419), eluted in water, and desalted with Centrisep columns (Princeton Separations cat. # CS-901). The purified mRNA-peptide fusion was reverse transcribed with Superscript IV (Invitrogen cat. # 18091050).

The purified library of peptide-cDNA conjugates was then incubated with biotinylated, photocleaved PhoCl and immobilized on streptavidin or neutravidin beads (Thermo Fisher Scientific, cat. # 29200, 20353) in selection buffer (20 mM Tris-HCl, pH 8.0, 150 mM NaCl, 0.02% Tween-20, 0.5 mg/ml BSA, 1 mM DTT, 1% FBS, and 0.2 mM D-Biotin). Full-length PhoCl was added to the reaction in excess. Incubation of the library with cleaved PhoCl for rounds 1 and 2 was conducted at 4°C, rounds 3-5 at room temperature, and round 6 at 30°C. The beads were then washed three times with selection buffer and once with TBST (20 mM Tris-HCl, pH 8.0, 150 mM NaCl, 0.02% Tween-20). The beads containing cleaved PhoCl and any peptides bound were used directly as template for PCR amplification of the library. The resulting PCR product was used as the library for the subsequent round of selection. The library underwent 6 rounds of selection.

### Ubiquitination and proteasome dependency of PFE3

COS-7 cells (ATCC cat# CRL-1651) between P5 and P20 were cultured in Dulbecco’s Modified Eagle’s Medium (DMEM) (ATCC, Cat # 30-2002) supplemented with 10% FBS (Hyclone, cat # SH30071.03) and 12.5mg/L gentamicin solution (Thermo Fisher Scientific, cat. # 15750-078) in a 5% CO_2_ incubator at 37°C. Cells were grown to 50% confluency and co-transfected with PSD-95.myc, reverse tetracycline transactivator (rtTA) and doxycycline-inducible PSD-95.FingR-Mdm2.RING or Rand.FingR-Mdm2.RING using Lipofectamine 2000 transfection reagent (Thermo Fisher Scientific, cat # 11668019).

24 hr after transfection, cells were treated with 1 µg/ml doxycycline (Sigma Aldrich, cat. # 5207) to induce expression of PSD-95.FingR-Mdm2.RING and 20 µM TAK243 (MedChemExpres, cat. # HY-100487) to block ubiquitination for 4 hr. Cells were then rinsed with PBS and scraped in Lysis Buffer (150 mM NaCl, 1 mM EDTA, 20 mM Tris, pH 8.0, 1% NP-40) with freshly added complete mini protease inhibitor cocktail (Roche, cat. # 04693124001). After incubating on ice for 30 min, the total cell lysate was centrifuged at 10,000rpm for 10 min at 4°C to pellet the insoluble fraction of the lysate. The concentration of the lysate was determined with a Bradford Protein Assay (BioRad, cat. # 5000201). 30 µg of cell lysate was denatured in Laemmli Buffer (BioRad, cat. # 1610747) with 10% 2-mercaptoethanol and ran on precast Any kD SDS-PAGE protein gels (BioRad, cat. # 4569034). Proteins were then transferred onto low-fluorescence PVDF membrane (BioRad cat. # 1620264) and probed with chicken anti-myc (1:3000, Novus Biologicals NB600-334) and mouse anti-α-tubulin (1:5000, Sigma T6199). Secondary antibodies were goat anti-chicken Alexa Fluor 680 (1:1000, Abcam, cat. # ab175779) and goat anti-mouse Alexa Fluor 750 (1:1000, Thermo Fisher Scientific, cat. # A-21037).

Blots were imaged with the Odyssey Infrared Imaging System (LI-COR Biosciences), and protein levels were determined using ImageJ Software (US National Institutes of Health). PSD-95 levels were normalized to α-tubulin levels.

### COS7 cell culture and transfection

COS-7 cells (ATCC cat# CRL-1651) were cultured in Dulbecco’s Modified Eagle’s Medium (DMEM) (ATCC, Cat # 30-2002) supplemented with 10% FBS (Hyclone, cat # SH30071.03) and 12.5 mg/L gentamicin solution (Thermo Fisher Scientific, cat. # 15750-078) in a 5% CO_2_ incubator at 37°C. Cells were grown to 50% confluence on coverslips (VWR, cat # 48393-059) and co-transfected Golgi-targeted PhoCl and tagRFP or PBP-tagRFP using Lipofectamine 2000 transfection reagent (Thermo Fisher Scientific, cat # 11668019). 24 hr after transfections, cells were transferred to Imaging Buffer.

### Excitatory synapse ablation with PFE3

Neurons for PSD-95 ablation were transfected as described above with 1µg pCAG:Zreg:tagRFP-PSD-95.FingR, 500ng CMV:Tet3G, and 1µg TRE-PFE3-HA or 1ug TRE-PFE3-P2A-GFP and incubated for four days. Neurons were then transferred to Imaging Buffer for imaging and positions of each neuron were saved in the Metamorph Software. Neurons were then transferred back into the cultured media and Doxycycline was added to 1µg/ml. After incubation for 48 hours, the same neurons were imaged to track changes in PSD-95.FingR. To confirm loss of endogenous PSD-95, neurons were immediately fixed and immunostained after imaging. To recover PSD-95 puncta, neurons were washed with conditioned media 5 times to remove the doxycycline and incubated in conditioned media for 5 days.

### Photocleavage of PhoCl in live cells

Neurons for synapse ablation were transfected with 1µg pCAG:tagRFP-GPHN.FingR-PhoCl2c and 500 ng pCAG:PBP-E3 and incubated for 5 days. COS7 cells were transfected with 200ng pGW:GTS-PhoCl2c and 50ng pCAG:PBP-tagRFP and incubated overnight. The COS7 cells and neurons were then imaged in Imaging Buffer and positions for each cell were recorded and saved. Each cell was subsequently illuminated with ∼400 nm light at 40mW/cm^2^ for 10 seconds every 30 seconds for a total of 3 minutes (total 1 min illumination time). The cells were then transferred to media and incubated at 37°C. Following the incubation, cells were again transferred to Imaging Buffer and imaged.

### PBP binding to cleaved PhoCl

PhoCl2c was expressed in HEK293T cells (ATCC Cat # CRL-3216) cultured in Dulbecco’s Modified Eagle’s Medium (DMEM) (ATCC, Cat # 30-2002) supplemented with 10% FBS (Hyclone, cat # SH30071.03), 125 mg/L gentamicin solution (Thermo Fisher Scientific, cat. # 15750-078) and GlutaMAX Supplement (Thermo Fisher Scientific, cat # 35050061) in a 5% CO_2_ incubator at 37°C. Cells were grown to 75% confluency in 150 mm cell culture dishes (Corning, cat # 430599) and transfected using PEI (Polysciences, cat # 24765-1) with Myc-PhoCl-GST or Myc-PhoCl-HRV3C.cleavage site-GST. After 72 hr of expression, cells were rinsed with PBS and lysed with ice cold Lysis Buffer (25mM Tris-HCl pH 7.4, 150 mM NaCl, 1 mM EDTA, 1% NP-40, 5% Glycerol) supplemented with freshly added complete mini protease inhibitor cocktail (Roche, cat. # 04693124001). The total cell lysate was centrifuged at 5000 rpm for 30 min at 4°C and the supernatant was bound to Glutathione Agarose beads (Thermo Fisher Scientific, cat # 16100) at 4°C overnight. The beads were then washed 5 times with Wash Buffer 5 (25 mM Tris-HCl pH 7.4,150 mM NaCl, 1 mM EDTA, 5% Glycerol). To generate photocleaved PhoCl2c, beads with Myc-PhoCl-GST were transferred to glass chambers (Cellvis, cat # C2-1.5H-N) and exposed to 400 nm LED 100 mW/cm^2^ for 20 seconds every minute for a total of 5 minutes. The beads were then transferred to an ultracentrifuge tube and rotated at 4°C overnight in Lysis Buffer to allow dissociation of PhoCl2c. To generate full-length PhoCl2c, beads containing Myc-PhoCl2c-HRV3C.cleavage site-GST were incubated with HRV3C protease (Thermo Fisher Scientific, cat # 88947) at 4°C overnight. The supernatant containing purified photocleaved or full-length PhoCl2c was separated from the beads using Spin-X centrifuge tube filters (Corning, cat # CLS8162). Photocleavage and HRV3C cleavage of photocleaved and full-length PhoCl2c was confirmed with SDS PAGE.

### Electrophysiology

To express channelrhodopsin-2 in type 6 cone bipolar cells, we crossed a CCK-Cre mouse line (Jackson Laboratory strain 012706) with the Jackson Laboratory Ai27 mouse, which contains a floxed channelrhodopsin2-TdTomato fusion protein sequence. Mice were intravitreally injected with either the PFE3 or control virus (1.5 µl). After 3-4 weeks, retinas were isolated and mounted in a recording chamber for patch clamp recording, as described previously (Jones et al., 2012). Retinas were perfused in ACSF bubbled with 95%O_2_/5%CO_2_. ACET (1 µM) and L-AP4 (10 µM) were added to block synaptic transmission from photoreceptors to Off and On bipolar cells, respectively. Infected cells were identified by GFP expression. GFP-positive On a-RGCs were identified by their large cell bodies, and their identity was confirmed by dye filling. Full-field 495 nm light (0.9 mW/cm^2^) was delivered to the retina for 10 ms to trigger EPSCs. Cells were voltage-clamped at −60 mV.

### Chemogenetic GFE3

Cultured cortical neurons were prepared and transfected with 1µg tagRFP-GPHN.FingR-HaloTag and 250ng eDHFR-E3 as described above at 14DIV. Neurons were imaged at 18DIV in Imaging Buffer and coordinates of each neuron were saved in the Metamorph Software. Immediately after imaging, neurons were transferred back to the culture media and TH was added to the medium at 100nM final concentration. After 4 or 24 hours of incubation, neurons were transferred to Imaging Buffer and the same neurons were imaged to track changes in GPHN.FingR. To confirm loss of endogenous Gephyrin, neurons were immediately fixed and immunostained as described above. To recover inhibitory synapses, the cultured neurons were washed three times with conditioned media and incubated with conditioned media with 50mM TMP for 48 hrs.

