## Supplementary Figures for "A toolbox for ablating excitatory and inhibitory synapses"

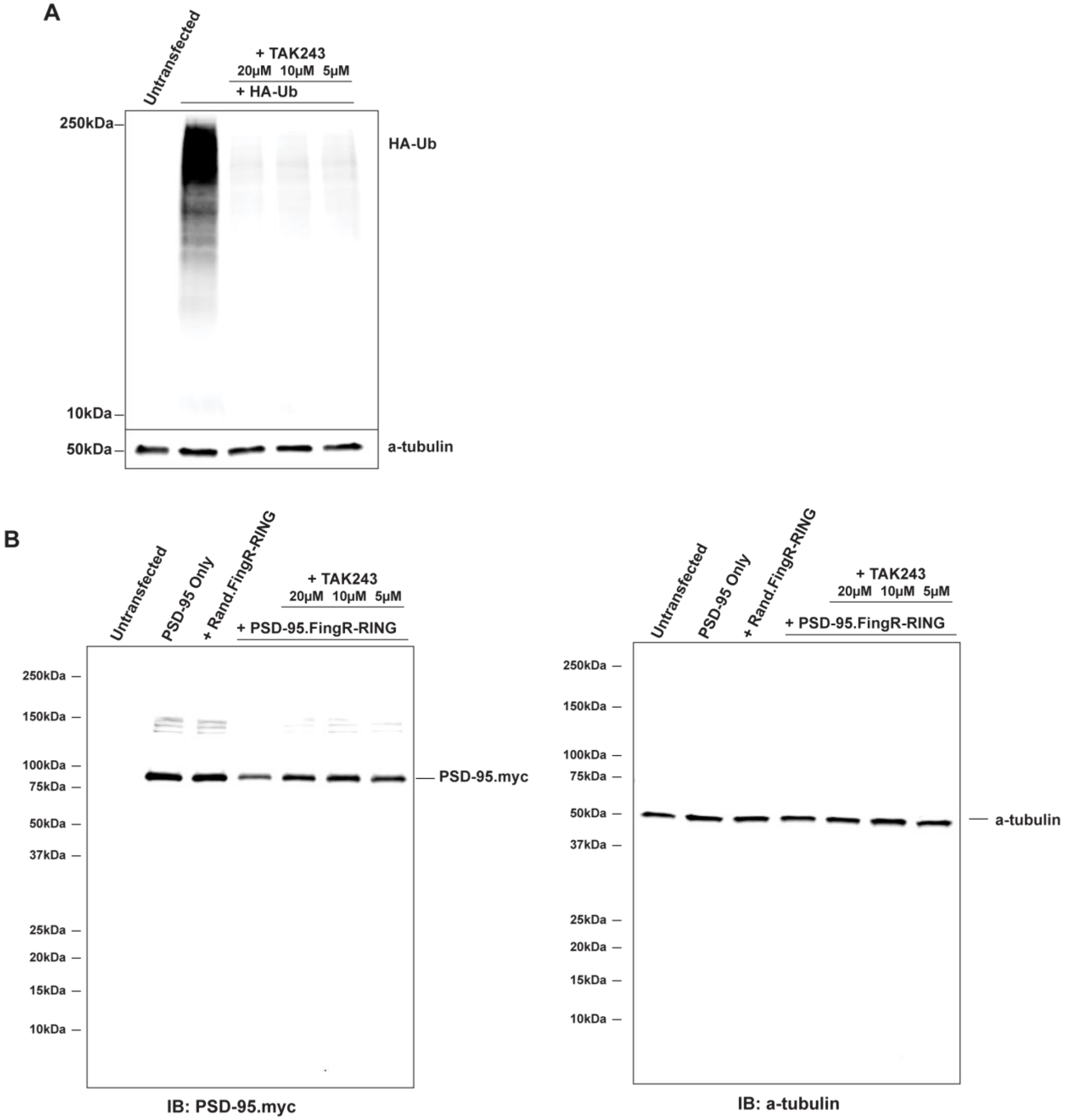


**Figure S1**: **TAK243 blocks ubiquitination and degradation of PSD-95 by Mdm2.RING.**

**A)** Western blot of lysate from COS7 cells transfected with HA-ubiquitin treated with TAK243 for 4 hours. Addition of TAK243 blocks high molecular weight bands indicating ubiquitination of proteins.

**B)** Full blot from Fig 1A.

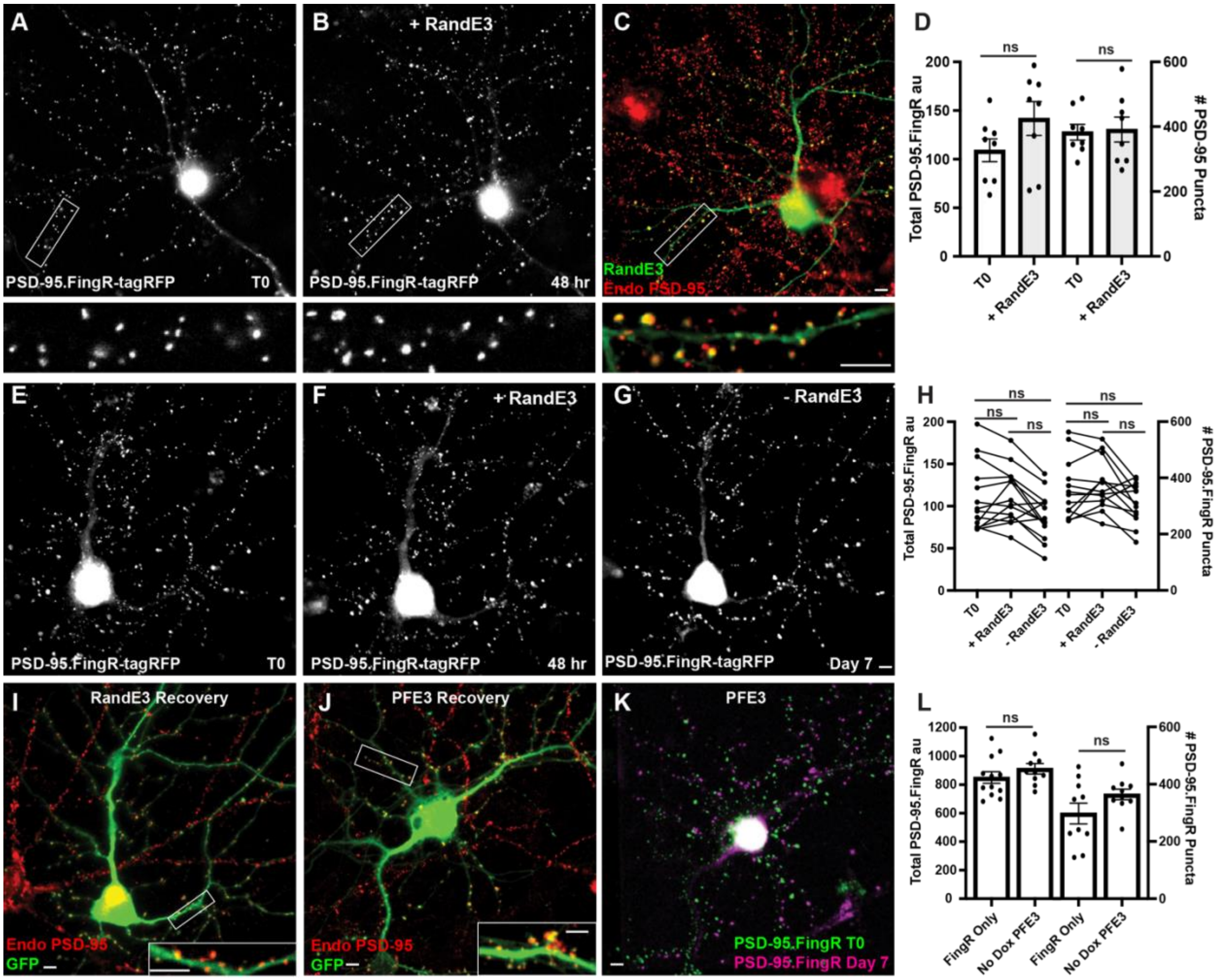


**Figure S2**: Expression of RandE3 does not significantly affect PSD-95 expression.

**A)** Cultured cortical neuron expressing PSD-95-FingR-tagRFP. Closeup of the boxed area below.

**B)** Same neuron as in A) after induction of RandE3 showing no apparent reduction in PSD-95-FingR-tagFRP labeling. Closeup of the boxed area below.

**C)** Neuron in B immunostained for RandE3 in green and endogenous PSD-95 in red. Closeup of the boxed area below shows the persistence of PSD-95 labeling.

**D)** Quantifications of the total amount of PSD-95.FingR-tagRFP labeling and the # of PSD-95.FingR puncta before and after expressing RandE3 show no significant changes in either measure. ns, $p > 0.05$, Kruskal-Wallis multiple comparisons.

**E)** Cultured cortical neuron expressing PSD-95-FingR-tagRFP.

**F)** Same neuron as in E) 48 hours after expressing RandE3 shows no apparent reduction in PSD-95-FingR-tagFRP labeling.

**G)** Same neuron as in E), F) 5 days after removal of Dox shows no apparent reduction in PSD-95-FingR-tagFRP labeling.

**H)** Quantifications of the total amount of PSD-95.FingR labeling and the # of PSD-95.FingR puncta indicate no significant changes with the addition and removal of Dox. Ns $p > 0.05$

**I)** Immunocytochemistry of neuron in G showing the neuron subjected to expression and removal of RandE3-P2A-GFP showing GFP in green and endogenous PSD-95 in red.

**J)** Immunostaining for neuron in Fig. 2K subjected to expression and removal of PFE3-P2A-GFP showing GFP in green and endogenous PSD-95 in red as in I). PSD-95 is present on the tips of dendritic spines.

**K)** Overlay of PSD-95.Fing-tagRFP labeling at T0 before expressing PFE3 (green) and after recovery of synapses at 7 days (purple) for the neuron shown in J) confirms the two labeling distributions are similar.

**L**) Total PSD-95.FingR labeling and number of puncta labeled with PSD-95.FingR in cultures expressing PSD-95.FingR alone vs. those expressing PFE3 without Dox are not significantly different ($P > 0.2$. Mann Whitney for total labeling, $P > 0.4$ Mann Whitney for # of puncta).

Scale bars represent 5 µm.

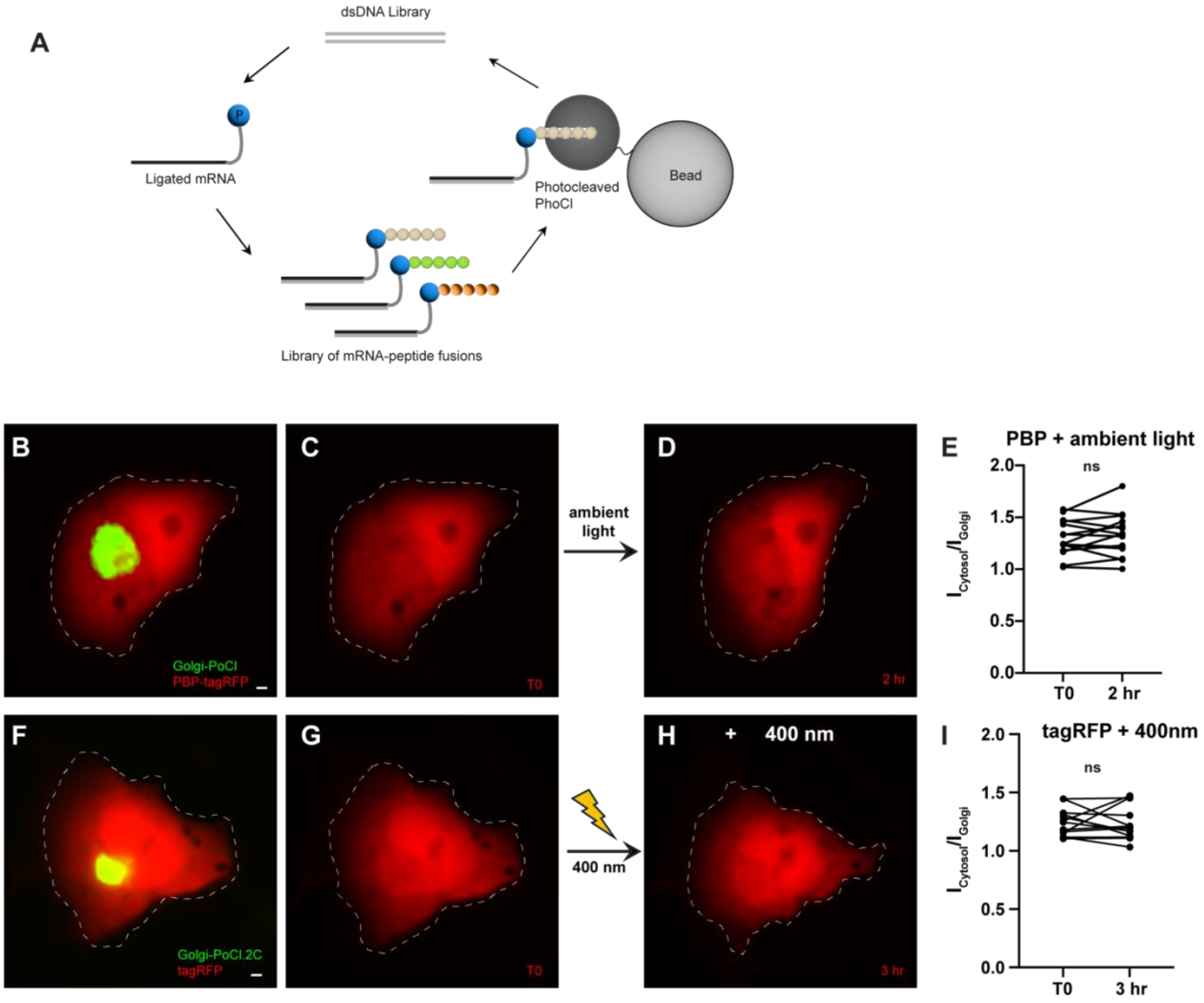


**Fig S3**: PhLIC does not activate with ambient light.

**A)** mRNA Display: The library of mRNA-protein fusions is incubated with cleaved PhoCl immobilized on beads. Peptides that bind cleaved PhoCl are pulled down, and the genetic material is amplified with PCR. The PCR product is transcribed, translated, and used to form mRNA-protein fusions, which are used for the subsequent round of selection.

**B), C)** COS7 cell expressing Golgi-targeted (PhoCl-GTS) and PBP-tagRFP before exposure to ambient light.

**D)** Same cell as in B), C) 2 hours after exposure to ambient light. There is no apparent targeting of PBP-tagRFP to the Golgi.

**E)** $I_{cytosol}/I_{Golgi}$ for PBP-tagRFP is not significantly different before vs. after exposure of cells to ambient light (+2 ± 2%, $p > 0.5$, Wilcoxon, $n = 15$ cells, 2 experiments).

**F), G)** COS7 cell expressing Golgi-targeted PhoCl2c-GTS and tagRFP. There is no apparent targeting of tagRFP to the Golgi.

**H)** Same cell as in F), G) 3 hours after illumination with 400 nm. There is no apparent targeting of PBP-tagRFP to the Golgi.

**I)** $I_{cytosol}/I_{Golgi}$ for tagRFP before and after illumination with 400 nm light does not differ ($0 \pm 3$ %, $p > 0.7$, Wilcoxon, $n = 13$). Scale bars represent 5 µm.

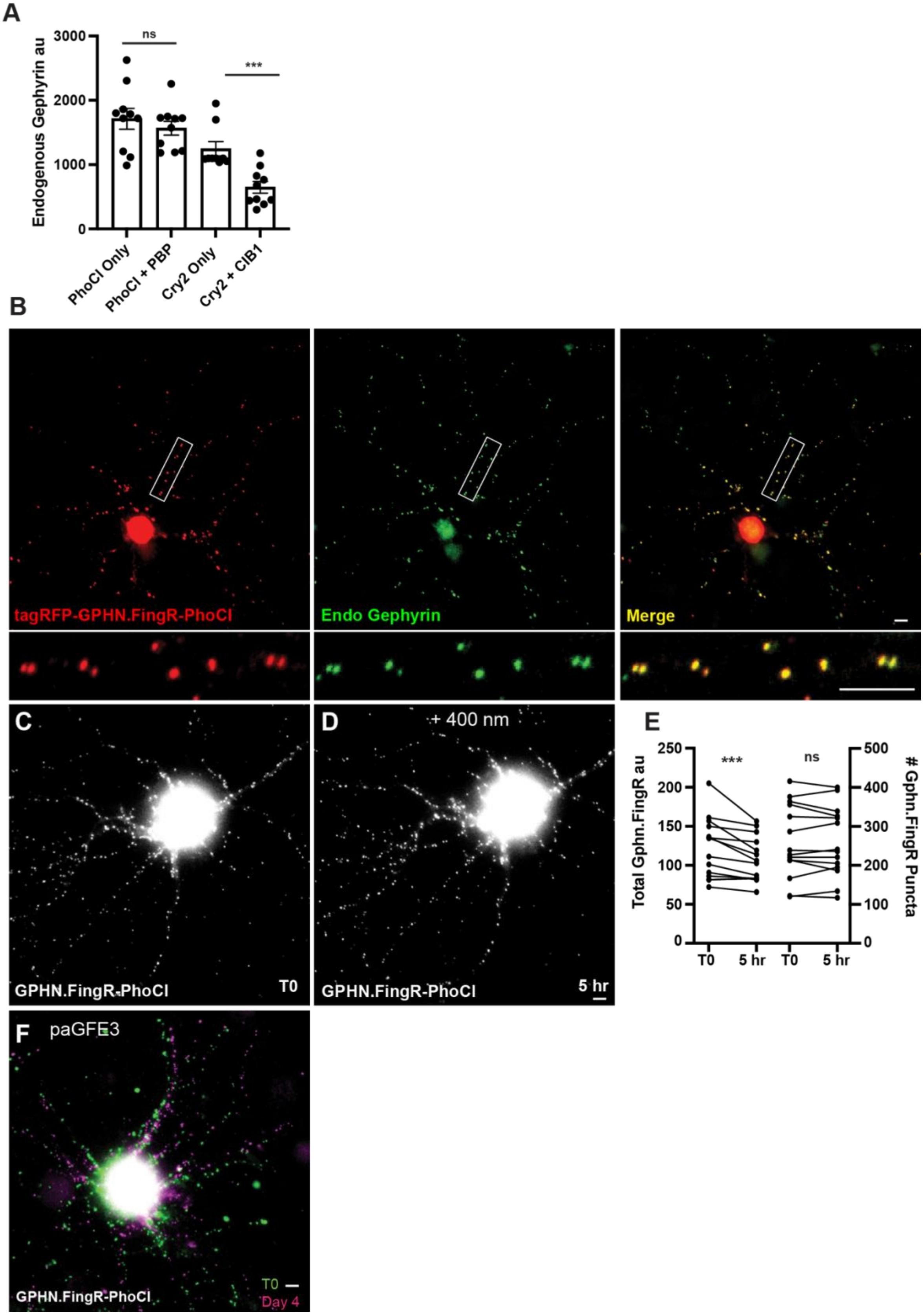

A
Endogenous Gephyrin au
3000
2000
1000
0
ns
***
PhoCl Only
PhoCl + PBP
Cry2 Only
Cry2 + CIB1
B
tagRFP-GPHN.FingR-PhoCl
Endo Gephyrin
Merge
C
GPHN.FingR-PhoCl
T0
D
+ 400 nm
GPHN.FingR-PhoCl
5 hr
E
Total Gphn.FingR au
250
200
150
100
50
0
***
ns
### Gphn.FingR Puncta
500
400
300
200
100
0
T0
5 hr
T0
5 hr
F
paGFE3
GPHN.FingR-PhoCl
T0
Day 4

**Fig S4 pa-GFE3 has no background activity**.

**A)** Background activity for the Cryptochrome (Cry2) photoactivation system and PhLIC 5 days after transfection. Cultured neurons transfected with GPHN.FingR-PhoCl + PBP-E3 did not show a significant difference in the intensity of staining with GPHN.FingR-tagRFP compared with those transfected with GPHN.FingR-PhoCl2c only. However, there was a significant reduction in endogenous Gephyrin between cultured neurons transfected with Gephyrin.FingR-Cry2 alone vs. with CIB1-E3.

**B)** Cultured neuron expressing transcriptionally regulated tagRFP-Gephyrin.FingR-PhoCl immunostained for tagRFP (red) and endogenous Gephyrin (green). Merge shows colocalization of tagRFP-Gephyrin.FingR-PhoCl with endogenous Gephyrin in yellow.

**C)** Neuron expressing GPHN.FingR-PhoCl2c alone before exposure to 400 nm light.

**D)** Same neuron as in C) after exposure to 400 nm light. There is no apparent loss of labeling.

**E)** Quantitation of labeling by tagRFP-GPHN.FingR-PhoCl2c in cells transfected with GPHN.FingR-tagRFP alone and then exposed to 400 nm for 1s and incubated for 5 hrs. Staining with GPHN.FingR-tagRFP was compared before and after light exposure. *** $p < 0.001$, ns $p > 0.05$

**F**) Same neuron as in Fig. 5F-H showing a comparison of tagRFP-GPHN.FingR-labeled puncta before (green) vs. after (magenta) exposure to 400 nm light.

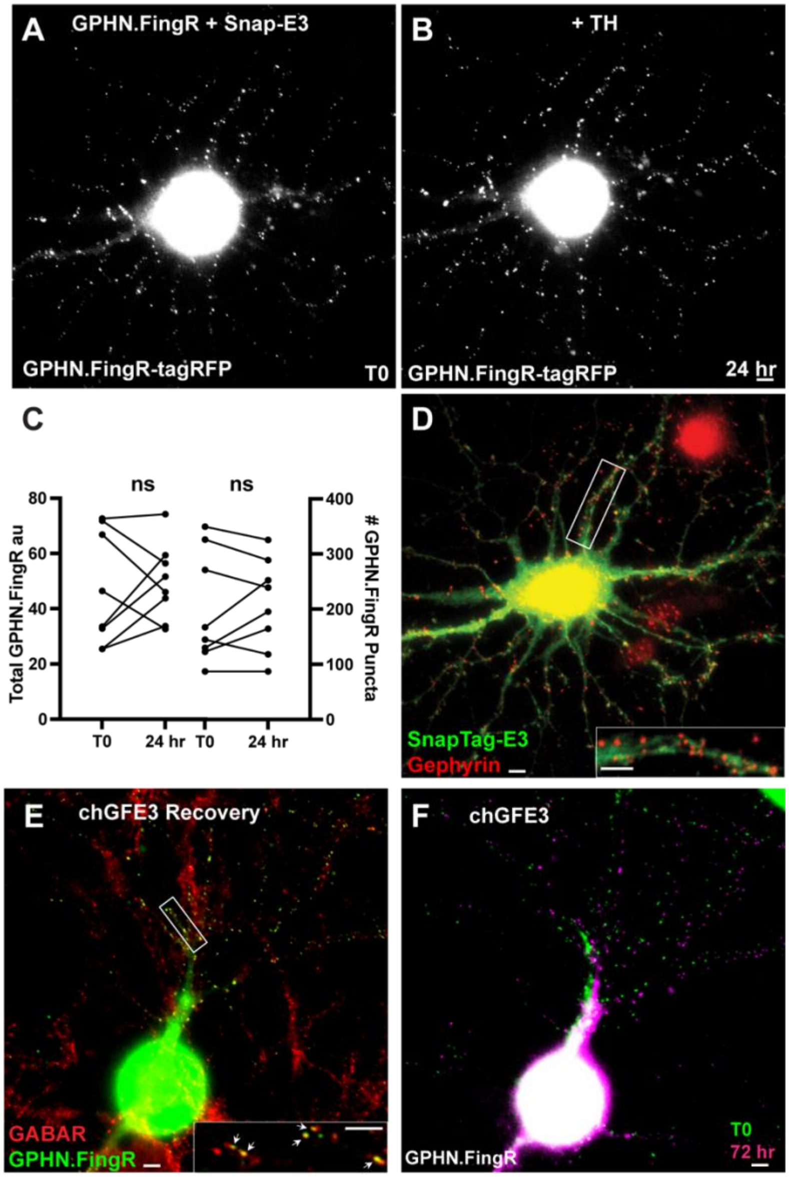


**Fig. S5** Chemogenetic ablation of inhibitory synapses with chGFE3

**A)** Cultured cortical neuron expressing tagRFP-Gephyrin.FingR-HaloTag and Snap-E3 for 4 days.

**B)** Same neuron as in A) 24 hours after addition of 100nm TH

**C)** Quantification of total Gephyrin and number of Gephryin puncta labeled by the GPHN.FingR-HaloTag. Incubation of the neurons co-expressing tagRFP-Gephyrin.FingR-HaloTag and Snap-E3 with TH did not affect Gephyrin (ns, $p > 0.05$, Wilcoxon).

**D)** Immunocytochemistry of the neuron in B) for endogenous Gephyrin (red) and Snap-E3 (green). Inset shows closeup of boxed area.

**E)** Immunocytochemistry of the neuron in Fig. 6M for $GABA_A$ receptors in red and GPHN.FingR in green. Closeup of the boxed area shows colocalization of the GPHN.FingR and $GABA_A$ receptor (arrows).

**F)** Overlay of the GPHN.FingR at T0 before the addition of TH (Fig. 6K) and after the addition of TMP to recover synapses at 72 hr (Fig. 6M).

Scale bars represent 5 µm
